# *In situ* loading of biologics on and tissue redirection of circulating extracellular vesicles with an anchoring nanobody

**DOI:** 10.64898/2026.10.26.761597

**Authors:** Gang Han, Li Zhong, Huanan Yu, Rongyan He, Yi Lu, Biaobiao Wang, Boyue Wu, Tongyang Zou, Xi Chen, Jingqiao Wu, Yao Zhang, Toshifumi Yokota, Yiqi Seow, HaiFang Yin

## Abstract

Extracellular vesicles (EVs) are promising nanocarriers of therapeutic macromolecules, but *ex vivo* production, cargo loading and quality control constrains clinical translation. Here we demonstrate that N_EV3_, an EV-anchoring nanobody identified by phage display, binds EVs from multiple sources through interactions involving fibronectin and loads diverse proteins onto EVs, resulting in enhanced intracellular delivery and biological activities *in vitro* and *in vivo*. Conjugating N_EV3_ to target proteins hijacks circulating EVs, prolonging circulation and altering tissue distribution *in vivo*. When fused with an GFP nanobody or protein A, N_EV3_ enabled simultaneous capture and loading of circulating GFP or immunoglobulin G (IgG) antibodies on endogenous EVs with enhanced intracellular delivery *in vitro* and redistribution to liver *in vivo*. Intravenous protein A-N_EV3_ significantly reduced circulating IgG and anti-dsDNA antibodies in systemic lupus erythematosus mice with improved survival and reduced pathologies. Intravenous anti-transferrin receptor-protein A-N_EV3_ redirected circulating EVs to muscle and alleviated muscle pathologies in dystrophin-deficient *mdx* mice. Our study demonstrates that N_EV3_ enables *in situ* loading of protein therapeutics and tissue-specific delivery of circulating EVs, providing a modular strategy for loading cargo onto and redirecting circulating EVs *in vivo*.

## Introduction

Extracellular vesicles (EVs) are natural biological nanocarriers of macromolecules and can be engineered for delivery of biotherapeutics endogenously or exogenously^1,2^. Genetic modification of EV-producing cells^3–6^ or *ex vivo* manipulation of isolated EVs^7,8^ can achieve efficient cargo loading, but these approaches face challenges in manufacturing, batch-to-batch characterization, scale-up and regulatory control^9^. Therefore, strategies that recruit patients’ circulating EVs in the recipient and functionalize them *in situ* can separate the therapeutic construct from the purification of exogenous EVs and could potentially circumvent these hurdles.

Nanobodies are compact, single-domain antibody fragments that can be produced recombinantly and conjugated readily to functional proteins^10^ and nanobody-based protein therapeutics have shown promise in clinical use and clinical trials^11–13^. In addition, nanobodies have been used to display targeting ligands on EVs^14,15^ or to load different proteins on EVs by fusing cargo-binding nanobodies with EV scaffold proteins^16,17^. However, it remains unclear whether a soluble EV-binding nanobody can load biotherapeutics on endogenous circulating EVs after systemic administration and whether protein-binding and tissue-targeting modules can be applied simultaneously.

With this in mind, we performed phage display with our proprietary synthetic nanobody library against CP05-immobilized EVs^18^ and identified N_EV3_ as an EV-anchoring nanobody. N_EV3_ bound to EVs from different sources specifically via fibronectin. N_EV3_ acted as a modular adaptor for proteins, RNA-binding domains and genome-editing enzymes, and intravenously administered N_EV3_ constructs associated with circulating EVs *in vivo*. By coupling N_EV3_ to an anti-GFP nanobody^19^ or protein A^20^, we further demonstrated the simultaneous recruitment of circulating proteins and antibodies to endogenous EVs. Finally, we evaluated two therapeutic proof-of-concept applications - depletion of circulating immunoglobulin (Ig)G antibodies in systemic lupus erythematosus (SLE) mice, and anti-transferrin receptor 1 (TfR1) IgG antibody^21^-guided redirection of EVs to skeletal muscle in dystrophin-deficient *mdx* mice. Together, the experiments define a strategy for *in situ* EV loading with protein cargoes and this technology potentially circumvents existing technical and regulatory constraints associated with *ex vivo* manipulation by harnessing circulating EVs as nanocarriers through a coordinated delivery of EV-anchoring nanobodies with protein of interest as a single entity. This approach may shift EV-based delivery towards nanobody-enabled protein therapeutics.

## Results

### N_EV3_ efficiently binds EVs from multiple sources

To identify EV-anchoring nanobodies, we performed *in vitro* biopanning with a synthetic nanobody library bearing humanized framework region backbones against differentiated mouse myotube EVs immobilized with CP05 (Fig.1a and Supplementary Fig. 1a)^18^. After seven rounds of selection, N_EV3_ was the most abundant sequence among 20 sequenced candidates (Fig. 1b), and sequence alignment against the UniProt and NCBI databases identified no known motifs. When fused to mCherry, N_EV3_ showed a greater EV-associated fluorescence signal than N_EV11_ or the control GFP nanobody N_GFP_ (Fig. 1c). As orientation might affect the EV-binding ability, we placed mCherry either on the N or C terminus of N_EV3_, with more mCherry-positive EVs detected for mCherry-N_EV3_ than N_EV3_-mCherry across different ratios (Supplementary Fig. 1b), indicating that accessibility of the C terminus of N_EV3_ contributes to EV binding. Consistently, more mCherry-positive EVs were found for mCherry-N_EV3_ incubated with EVs from human mesenchymal stem cells, 293T and Panc02 cells, and mouse serum, dendritic cells and differentiated myotubes than mCherry-N_GFP_ (Fig. 1d and Supplementary Fig. 1c). Cy5-labeled N_EV3_ bound more WGA-AF488-labeled EVs from differentiated mouse myotubes than N_GFP_ (Fig. 1e), with an average of 135.9 Cy5-labeled N_EV3_ per vesicle detected by Nanoimager (Fig. 1f), though N_EV3_ loading did not measurably change EV morphology or size distribution (Supplementary Fig. 1a). Collectively, these results support the conclusion that N_EV3_ enables efficient EV binding and protein-loading irrespective of EV source.

**Figure 1.**
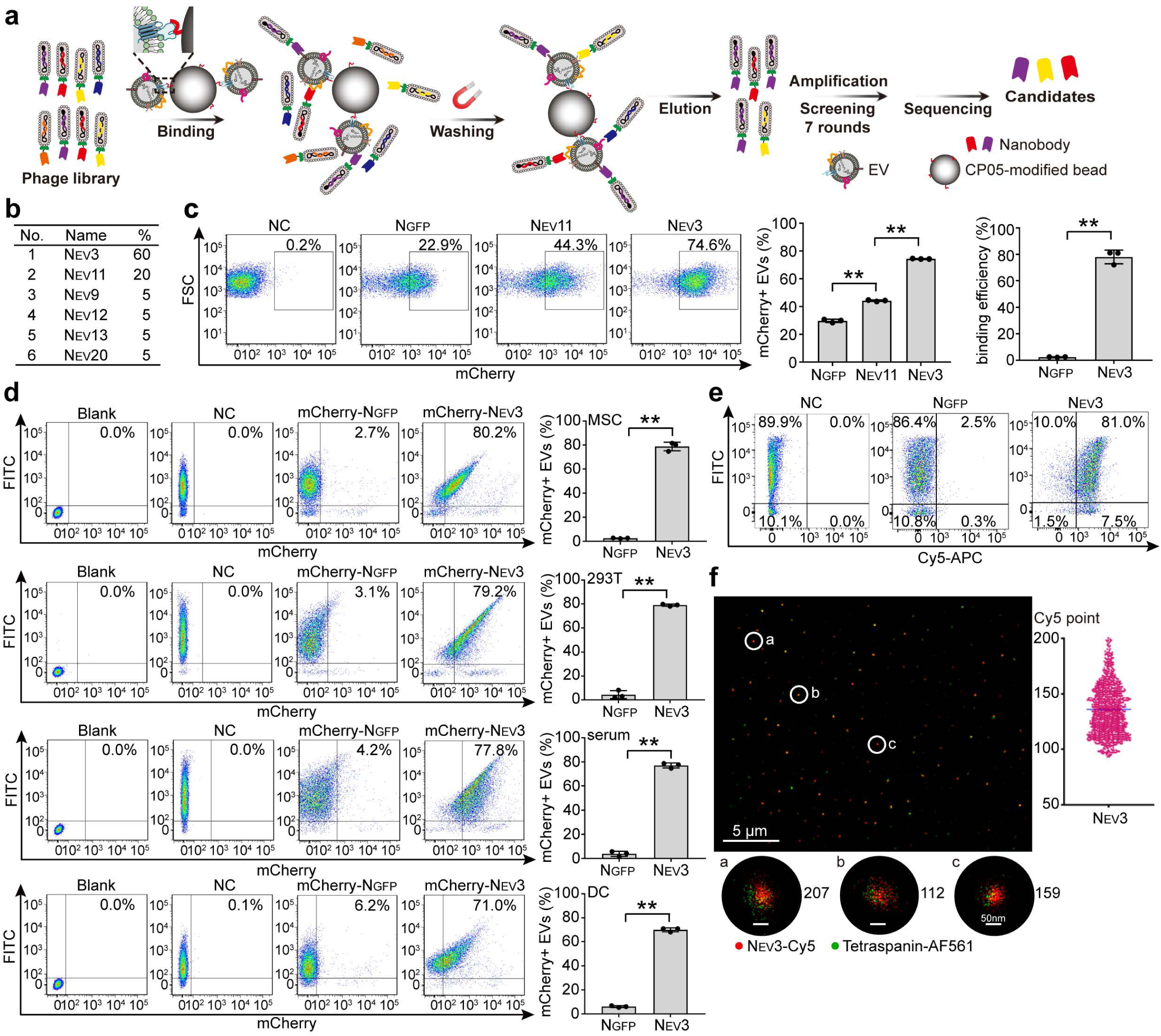
Screening and identification of the EV-anchoring nanobody N_EV3_. (**a**) Schematic of the screening strategy used to identify EV-anchoring nanobodies. (**b**) Table of top-ranked candidate nanobodies based on frequency. (**c**) Flow cytometry plots and quantification of mCherry-positive EVs after incubation of mCherry-N_EV3_, mCherry-N_EV11_ or mCherry-N_GFP_ with EVs derived from differentiated mouse myotubes (n=3; **p< 0.01; Welch’s one-way ANOVA followed by Holm-adjusted pairwise Welch’s t-tests). EVs alone were used as the blank control. (**d**) Flow cytometry plots and quantification of mCherry-positive EVs after incubation of mCherry-N_EV3_ or mCherry-N_GFP_ with EVs derived from human mesenchymal stem cells, 293T cells, mouse serum and mouse DC2.4 cells (n=3; **p< 0.01; two-sided unpaired Welch’s t-tests). EVs were labeled with WGA-AF488 and detected in the FITC channel. NC refers to WGA-AF488-labeled EVs; blank represents unlabeled EVs. (**e**) Flow cytometry plots and quantification of the binding of Cy5-labeled N_EV3_ or N_GFP_ to WGA-AF488-labeled EVs derived from differentiated mouse myotubes (n=3; **p< 0.01; two-sided unpaired Welch’s t-test). (**f**) Representative super-resolution microscopic images and quantification of Cy5-labeled N_EV3_ molecules associated with individual EVs derived from differentiated mouse myotubes. EVs were stained with AF561-conjugated antibodies against CD63, CD81 and CD9. Scale bars were 5 μm (overview) and 50 nm (magnified EVs), respectively.

### EV-associated fibronectin contributes to N_EV3_ binding

N_EV3_-coated beads recovered EV markers Alix and CD81^22^ from intact EVs derived from differentiated mouse myotubes, whereas N_GFP_-coated beads did not (Fig. 2a). To investigate whether the binding of N_EV3_ to EVs is protein-oriented, we blocked exposed EV proteins with gold nanoparticles (AuNPs)^23^, and subsequently incubated with mCherry-N_EV3_. The proportion of mCherry- and DiR-double-positive EVs declined with increasing concentrations of AuNPs and saturated at an EV-to-Au ratio of 1:20 (Fig. 2b). Digesting EV surface proteins with proteinase K reduced binding of mCherry-N_EV3_ to EVs from differentiated mouse myotubes (Supplementary Fig. 2a), whereas combined proteinase K and peptide-N-glycosidase F (PNGase F)^24^ treatment led to a larger reduction (Fig. 2c), indicating glycosylated proteins on the EV surface contribute substantially to N_EV3_ binding. Further immunoprecipitation revealed a series of protein bands with molecular weights above 170 kDa observed in the lane loaded with GST-tagged N_EV3_ and EVs but not in EVs alone (Supplementary Fig. 2b). Mass spectrometric analysis identified fibronectin, an extracellular matrix glycoprotein found on the surface of EVs^25,26^, as the most abundant candidate interactor based on label-free quantification (LFQ) intensity (Supplementary Fig. 2c), and fibronectin was recovered specifically with GST-tagged N_EV3_-coated beads (Fig. 2d). EVs generated from fibronectin-depleted cells showed less binding to N_EV3_ than control EVs (Fig. 2e and Supplementary Fig. 2d, e). Biolayer interferometry showed dramatically lower biological layer thickness for N_EV3_ incubated with fibronectin-depleted EVs than for control EVs, with the response decreasing from 0.2908 nm to 0.1379 nm (Fig. 2f), supporting a specific interaction between N_EV3_ and EV-associated fibronectin. AlphaFold3 modelling predicted interactions of N_EV3_ with seven functional domains of fibronectin, defined by Uniprot, with potential hydrogen bonds between E48, R57, R58, N59, K71, N80 and R108 of N_EV3_ and D745, V747, Q792, E795, S800, I1455, N1463, Q1534 and T1565 of fibronectin, respectively (Fig. 2g). Taken together, these data indicate that EV-associated fibronectin contributes substantially to N_EV3_’s binding.

**Figure 2.**
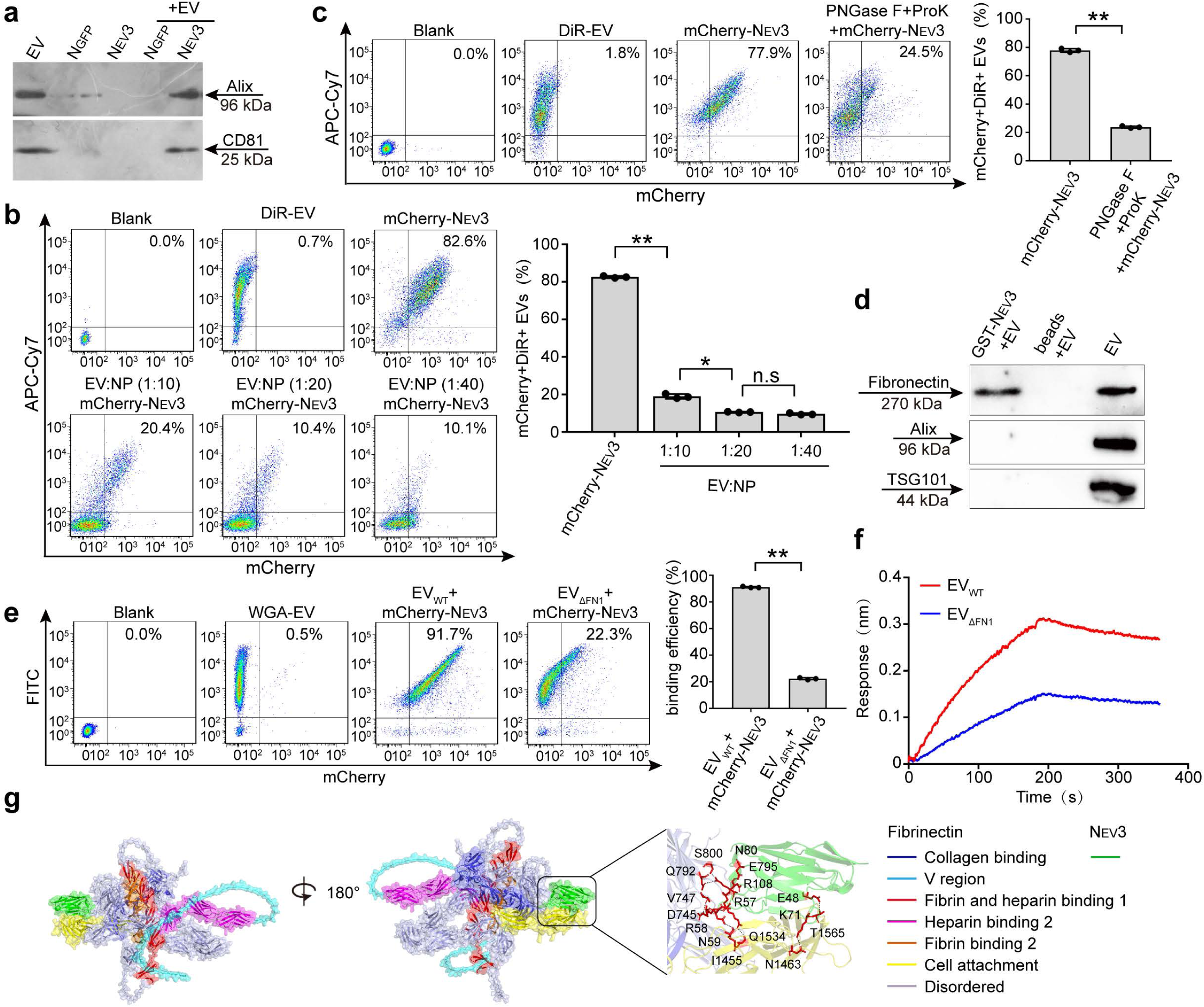
Identification and validation of N_EV3_’s binding partners on EVs. (**a**) Western blot analysis of EV markers Alix and CD81 in EVs captured by N_EV3_- or N_GFP_-coated beads. EVs were purified from differentiated C2C12 cells and resuspended in PBS. (**b**) Representative flow cytometry plots and quantification of mCherry- and DiR- positive EVs (n=3; Welch’s one-way ANOVA followed by Holm-adjusted Welch’s t- tests). EVs were incubated with AuNPs (NP) at the indicated EV-to-NP mass ratios, followed by co-incubation with mCherry-N_EV3_. EVs were derived from differentiated C2C12 cells. Blank refers to unlabeled EVs. (**c**) Representative flow cytometry plots and quantification of mCherry- and DiR-positive EVs (n=3; two-sided unpaired Welch’s t-test). EVs were digested with proteinase K (ProK) and PNGase F, followed by incubation with mCherry-N_EV3_. EVs were derived from differentiated C2C12 cells. (**d**) Western blot analysis of fibronectin in EVs pulled down by N_EV3_-GST beads. EVs were derived from differentiated C2C12 cells. EVs alone served as a positive control and beads incubated with EVs served as a negative control. (**e**) Representative flow cytometry plots and quantification of mCherry- and FITC-positive EVs (n=3; two-sided unpaired Welch’s t-test). mCherry-N_EV3_ was incubated with EVs derived from fibronectin-depleted 293T cells (EV_ΔFN1_) or 293T cells (EV_WT_), respectively. EVs were labeled with WGA-AF488. (**f**) Biolayer interferometry analysis of N_EV3_ binding to EV_WT_ and EV_ΔFN1_. (**g**) Predicted interaction between N_EV3_ and fibronectin using AlphaFold2. The predicted interacting residues and fibronectin domains are indicated. *p< 0.05; **p< 0.01. n.s. means not significant.

### N_EV3_ facilitates EV-associated delivery of macromolecules *in vitro* and *in vivo*

EVs loaded with mCherry-N_EV3_ were taken up by 293T cells more efficiently than were EVs incubated with mCherry-N_GFP_ as shown by a significantly higher proportion of mCherry- and AF488-double-positive cells (Fig. 3a). Given that intracellular delivery of small interfering RNA (siRNA) remains challenging, we investigated if siRNA delivery could be enhanced when loaded onto EVs via N_EV3_. To realize this, we fused N_EV3_ to a double-stranded RNA-binding domain (DRBD)^27^. *GAPDH* siRNAs^28^ bound to DRBD-N_EV3_ were loaded onto EVs, and EV-associated siRNA-DRBD-N_EV3_ showed increased uptake in 293T cells relative to siRNAs incubated with EVs alone (Fig. 3b, c and Supplementary Fig. 3). *GAPDH* expression was significantly reduced in cells treated with EV-associated siRNA-DRBD-N_EV3_ relative to siRNAs incubated with EVs alone (Fig. 3d). Accumulation of FITC signals was found in liver from mice intravenously injected with DiR-labeled EVs loaded with FITC-labeled siRNA-DRBD- N_EV3_ (Fig. 3e), supporting the conclusion that EVs facilitate siRNA delivery to liver via DRBD-N_EV3_. To investigate intranuclear delivery of protein therapeutics loaded on EVs via N_EV3_, we fused Cre recombinase with N_EV3_ and loaded it onto EVs with an efficiency of approximately 85.6% (Fig. 3f). EV-associated Cre-N_EV3_ induced a stark fluorescence switch from mCherry (red) to GFP (green) in Cre-loxP-based color-switching reporter cells and from tdTomato (red) to GFP (green) in transgenic mice^29^ after intramuscular injection, but not with Cre-N_GFP_ (Fig. 3g, h), confirming enhanced intranuclear delivery of Cre recombinase by EVs loaded via N_EV3_. Strikingly, N_EV3_ facilitated the loading Cas9 and Cas9/sgRNA ribonucleoprotein complex on EVs, with a loading efficiency of about 44.5% (Fig. 3i), resulting in enhanced intracellular delivery and genome-editing efficiency in CRISPR-Cas9 reporter cells compared with Cas9 incubated with EVs (Fig. 3j, k). These findings reinforce that N_EV3_ enables efficient loading of diverse protein therapeutics on EVs, resulting in improved intracellular and intranuclear delivery and biological activities *in vitro* and *in vivo*.

**Figure 3.**
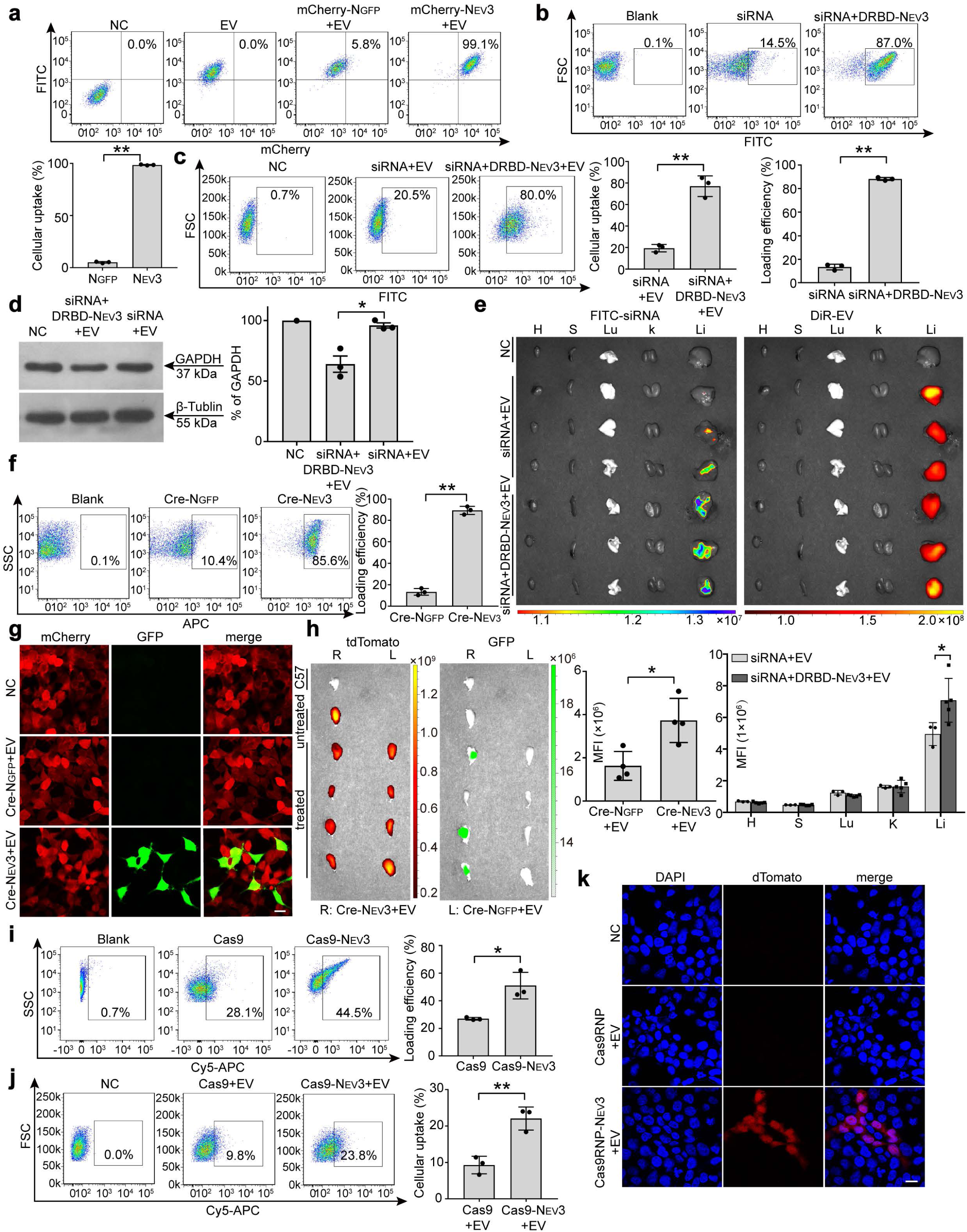
Evaluation of N_EV3_-mediated loading of different cargoes on exogenous EVs and biological activities *in vitro* and *in vivo*. (**a**) Representative flow cytometry plots and quantification of mCherry-positive cells (n=3; two-sided unpaired Welch’s t-test). Mixture of mCherry-N_EV3_ or mCherry-N_GFP_ with EVs, or EVs alone were incubated with 293T cells for 24 h prior to flow cytometry. EVs were derived from 293T cells and labeled with WGA-AF488. NC refers to cells treated with PBS. (**b**) Representative flow cytometry plots and quantification of FITC-positive EVs (n=3; two-sided unpaired Welch’s t-test). FITC-labeled siRNA-DRBD-N_EV3_ or FITC-labeled siRNA was mixed with EVs and examined by flow cytometry. EVs were derived from 293T cells. (**c**) Representative flow cytometry plots and quantification of FITC-positive cells (n=3; two-sided unpaired Welch’s t-test). EVs associated with siRNA-DRBD- N_EV3_ or a mixture of siRNA and EVs were co-incubated with 293T cells for 24 h and examined by flow cytometry. NC refers to cells treated with PBS. (**d**) Representative western blot and quantification of GAPDH in 293T cells treated with *GAPDH* siRNA-DRBD-N_EV3_-loaded EVs or a mixture of *GAPDH* siRNA and EVs for 48 h (n=3; two-sided paired t-test). β-tubulin was used as the loading control. A total of 50 μg protein was loaded per lane. (**e**) Tissue fluorescence images and quantification of FITC and DiR signals in body-wide tissues. MFI refers to mean fluorescence intensity unless otherwise specified. FITC-labeled siRNA-DRBD-N_EV3_-EVs (n=5) or a mixture of FITC-labeled siRNA and EVs (n=3) were intravenously injected into *C57BL/6* mice and tissues were harvested 1 h post-injection (two-sided unpaired Welch’s t-tests). EVs were derived from 293T cells and labeled with DiR. H, heart; S, spleen; Lu, lung; K, kidney; Li, liver; NC refers to mice injected with buffer alone. (**f**) Representative flow cytometry plots and quantification of Cy5-positive EVs (n=3; two-sided unpaired Welch’s t-test). Cy5-labeled Cre-N_EV3_ or Cre-N_GFP_ was incubated with EVs derived from 293T cells. (**g**) Representative fluorescence microscopy images of Cre-reporter 293T cells 72 h after treatment with Cre-N_EV3_-loaded EVs or a mixture of Cre-N_GFP_ and EVs (scale bar=10 μm). NC refers to cells treated with PBS. (**h**) Representative *ex vivo* fluorescence images and quantification of GFP signals in tibialis anterior (TA) muscles from Cre-reporter mice 7 days after intramuscular administration of Cre-N_EV3_-loaded EVs (right TA-R) or a mixture of Cre-N_GFP_ and EVs (left TA-L) (n=4; two-sided paired t-test). C57 refers to TA muscle from a wild-type *C57BL/6* mouse. (**i**) Representative flow cytometry plots and quantification of Cy5-positive EVs (n=3; two-sided unpaired Welch’s t-test). Cy5-labeled Cas9-N_EV3_ or Cas9 was incubated with EVs derived from 293T cells. (**j**) Representative flow cytometry plots and quantification of Cy5-positive cells (n=3; two-sided unpaired Welch’s t-test). Cy5-labeled Cas9-N_EV3_-loaded EVs or a mixture of Cas9 and EVs were co-incubated with 293T cells for 24 h prior to flow cytometry. NC refers to cells treated with PBS. (**k**) Representative fluorescence microscopy images of CRISPR-Cas9 reporter 293T cells after treatment with Cas9 RNP-N_EV3_-loaded EVs or a mixture of Cas9 RNP and EVs for 72 h (scale bar=10 μm). NC refers to cells treated with PBS. *p < 0.05; **p< 0.01.

### N_EV3_ prolongs and redistributes proteins by hijacking circulating EVs *in vivo*

We next investigated if N_EV3_ can capture circulating proteins and directly load proteins onto circulating EVs. To this end, we incubated N_EV3_-mCherry with cell supernatant or mouse plasma. The results showed that mCherry was detected in EV fractions from cell supernatant or mouse plasma but not with mCherry alone (Supplementary Fig. 4a, b). Corroborating with *in vitro* data, a substantial number of mCherry-positive EVs were recovered from blood of wild-type *C57BL/6* mice after intravenous injection of mCherry-N_EV3_ (Fig. 4a), with a longer half-life than mCherry-N_GFP_ (Fig. 4b). Concordantly, mCherry signals accumulated in liver from mice intravenously injected with mCherry-N_EV3_ compared with mCherry-N_GFP_ under identical conditions, though no difference was found in kidney (Fig. 4c). However, pharmacological reduction of circulating EVs with GW4869, an inhibitor of EV biogenesis and release^30^, decreased liver-associated mCherry signals and the signal was recovered after the addition of exogenous EVs (Supplementary Fig. 5a, b), strengthening the assumption that mCherry-N_EV3_ hijacks circulating EVs, resulting in redistribution of mCherry to liver. Significantly greater mCherry signals were found in liver sections and hepatocytes, macrophages and endothelial cells isolated from liver of mice intravenously injected with mCherry-N_EV3,_ than mCherry-N_GFP_ under identical conditions (Fig. 4d-f and Supplementary Fig. 6). Likewise, intravenous administration of R Rhodamine B - labeled siRNA-DRBD-N_EV3_ into wild-type *C57BL/6* mice resulted in significantly stronger fluorescence signals in liver than siRNA alone (Fig. 4g). Collectively, these findings demonstrate that N_EV3_ is capable of hijacking circulating EVs, resulting in prolonged circulation, redistribution and enhanced intracellular delivery of protein cargoes *in vivo*.

**Figure 4.**
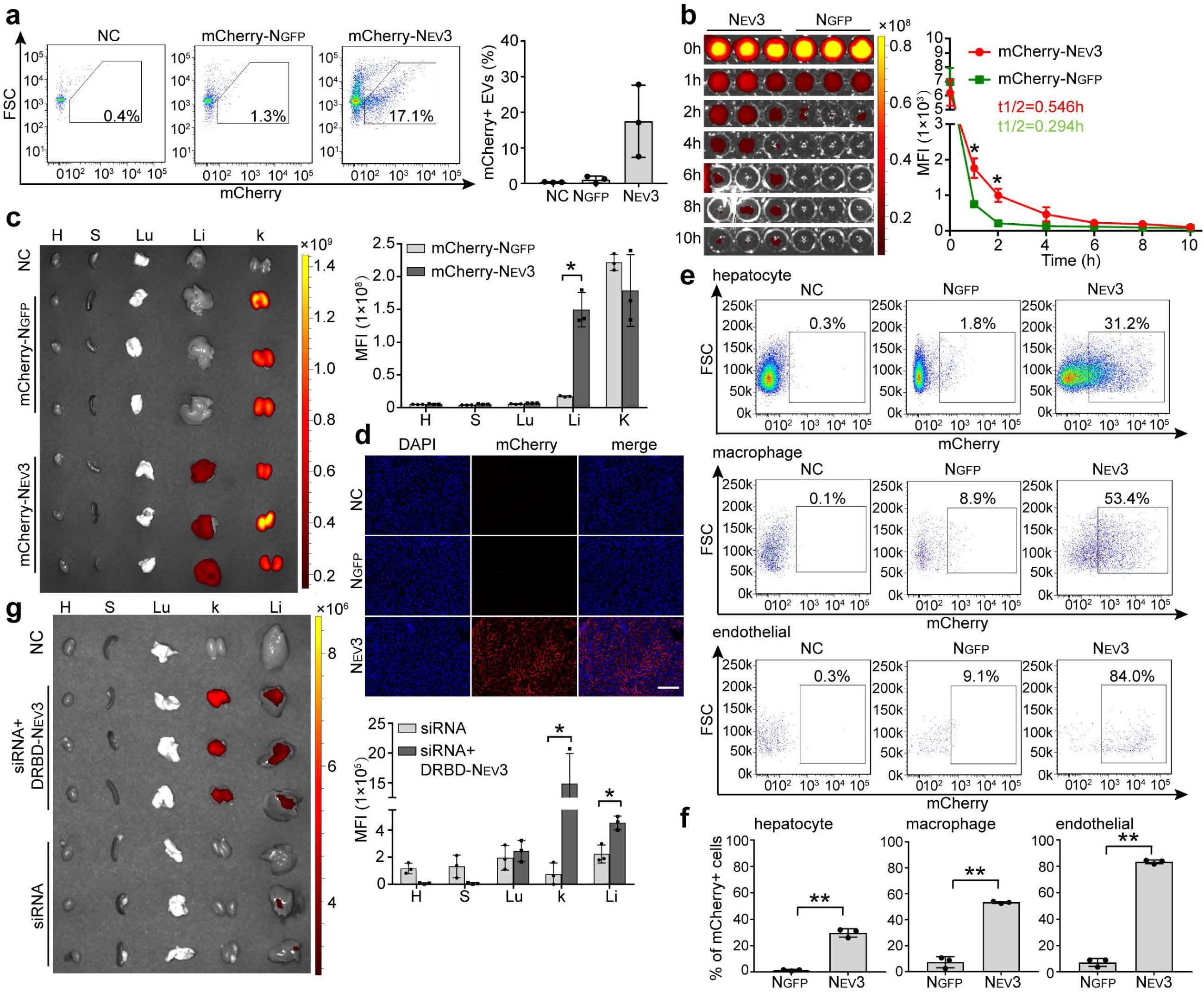
Evaluation of N_EV3_’s ability to capture circulating endogenous EVs *in vitro* and *in vivo*. (**a**) Representative flow cytometry plots and quantification of mCherry-positive EVs (n=3). mCherry-N_EV3_ or mCherry-N_GFP_ was injected into *C57BL/6* mice intravenously and plasma was collected 2 h post-injection, and EVs were isolated from plasma via centrifugation. NC refers to EVs isolated from mice injected with an equal volume of PBS. (**b**) Measurement of the plasma half-lives of mCherry- N_EV3_ and mCherry-N_GFP_ in *C57BL/6* mice (n=3). mCherry-N_EV3_ or mCherry-N_GFP_ was intravenously injected into *C57BL/6* mice and plasma was collected at 0, 1, 2, 4, 6, 8 and 10 h post-injection. Half-lives were calculated by GraphPad Prism. (**c**) Representative *ex vivo* fluorescence images and quantification of mCherry signals in body-wide tissues (n=3). mCherry-N_EV3_ or mCherry-N_GFP_ was intravenously injected into *C57BL/6* mice and tissues were harvested 2 h after injection. H, heart; S, spleen; Lu, lung; Li, liver; K, kidney; NC refers to mice injected with buffer. (**d**) Representative fluorescence microscopy images of liver sections from mice treated with mCherry-N_EV3_ or mCherry-N_GFP_ (scale bar=100 μm). Representative flow cytometry plots (**e**) and quantification (**f**) of mCherry-positive hepatocytes, macrophages and endothelial cells isolated from livers of *C57BL/6* mice injected with mCherry-N_EV3_ or mCherry-N_GFP_ (n=3). Livers were harvested 2 h after injection. NC refers to mice injected with buffer. (**g**) Representative *ex vivo* fluorescence images and quantification of intensity in body-wide tissues. Rhodamine B -labeled siRNA-DRBD-N_EV3_ or FITC-labeled siRNA was intravenously injected into *C57BL/6* mice and tissues were harvested 30 min after injection (n=3). H, heart; S, spleen; Lu, lung; K, kidney; Li, liver; NC refers to mice injected with buffer. Statistical significance was assessed by two-sided unpaired Welch’s t-tests; *p < 0.05; **p< 0.01.

### Modular N_EV3_ adaptors recruit circulating proteins to endogenous EVs

Having showcased the ability of N_EV3_ to hijack circulating EVs *in vivo*, we tested whether N_EV3_ can recruit circulating proteins to endogenous EVs. To this end, we fused N_EV3_ to the GFP-binding N_GFP_ (Fig. 5a). More GFP-positive EVs were detected when N_GFP_-N_EV3_, GFP and EVs were mixed (Fig. 5b), showing the ability of N_EV3_ to load circulating GFP on EVs. Concordantly, significantly increased GFP fluorescence signals were found in 293T cells (Fig. 5c) or in liver (Fig. 5d, e) when N_GFP_-N_EV3_, EVs and GFP were incubated with cells or sequentially injected into wild-type *C57BL/6* mice intravenously, compared with mixture of N_GFP_, EVs and GFP. These results further support N_EV3_’s ability to facilitate loading of circulating proteins on endogenous EVs, resulting in enhanced intracellular delivery and redistribution to liver by hitching a ride on EVs. Since protein A shows high binding affinity to fragment crystallizable (Fc) region of IgG antibodies^20^, we fused protein A to N_EV3_ and tested the recruitment of IgG antibodies onto EVs (Fig. 5f). Protein A-N_EV3_ facilitated the loading of anti-CD45 IgG antibodies on EVs, reflected by FITC and APC-Cy7 double-positive EVs when AF488-WGA-labeled EVs were mixed with APC-Cy7-anti-mouse CD45 IgG antibodies and protein A-N_EV3_ (Fig. 5g). Consistently, Protein A-N_EV3_ enabled simultaneous loading of four different IgG antibodies on EVs, exemplified by APC-Cy7-anti-mouse CD45, APC-anti-mouse CD80, BV421-anti-mouse XCR1, PE-Cy7-anti-mouse CD25 IgG and AF488-WGA-labeled EVs *in vitro* (Fig. 5h and Supplementary Fig. 7a). Protein A-N_EV3_ increased intracellular fluorescence while reduced the fluorescence remaining in the medium when protein A-N_EV3_, FITC-conjugated IgG antibodies and EVs were co-incubated with 293T or Hepa1-6 cells, compared with incubation with protein A-N_GFP_ (Fig. 5i and Supplementary Fig. 7b), supporting the conclusion that N_EV3_ facilitates loading of circulating IgG antibodies on EVs, resulting in enhanced intracellular delivery. Intravenous protein A-N_EV3_ at a dose of 2.5 mg/kg caused a transient reduction in circulating total IgG antibodies in wild-type *C57BL/6* mice but not with protein A-N_GFP_ under identical conditions (Fig. 5j), suggesting that protein A-N_EV3_-facilitated circulating EV-binding contributes to reduced levels of circulating IgG antibodies. Altogether, these data showed that N_EV3_ adaptor enables simultaneous capture of circulating protein and EVs, and loading of circulating proteins on endogenous EVs results in enhanced intracellular delivery *in vitro* and redistribution of proteins to liver *in vivo*.

**Figure 5.**
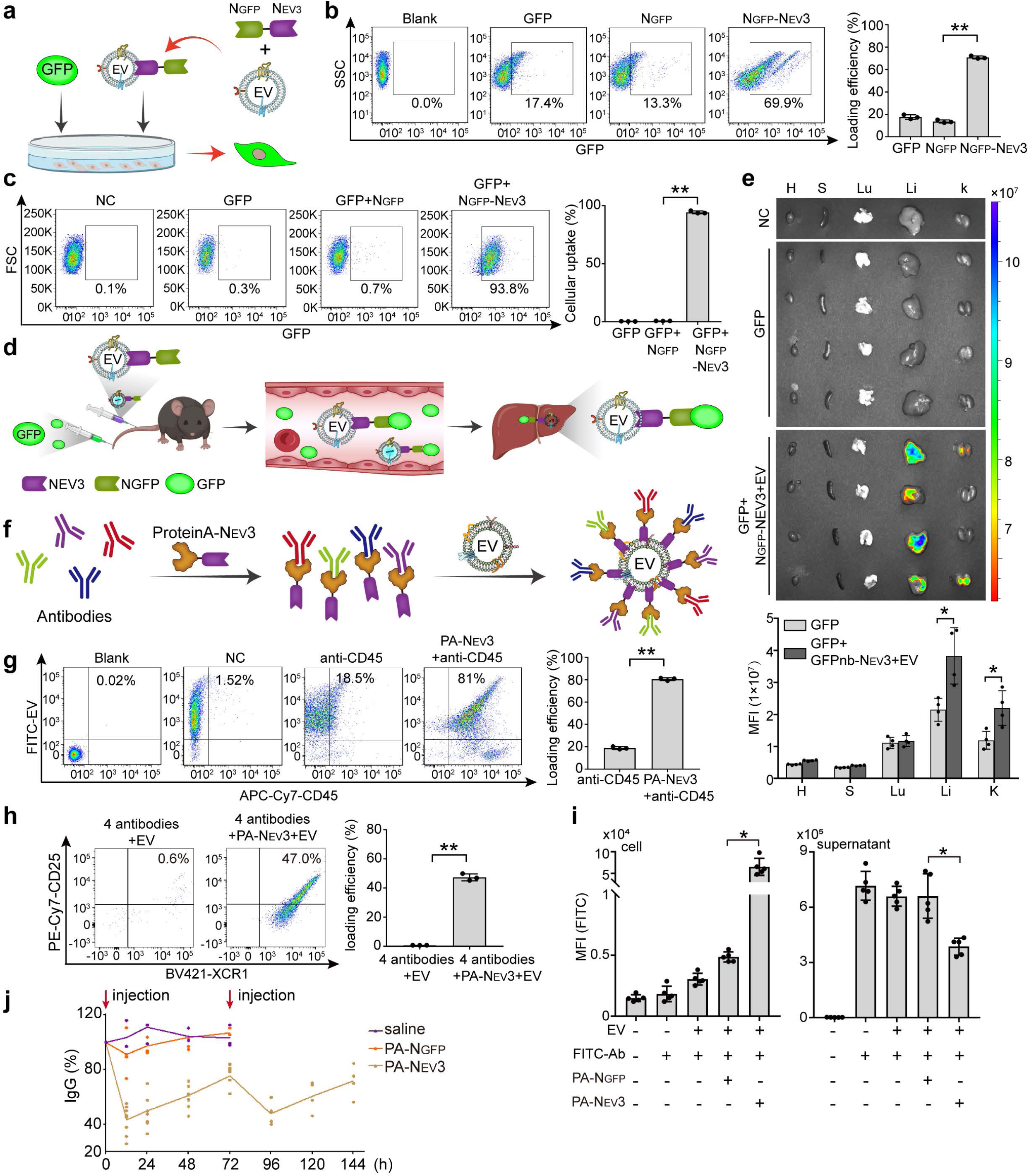
Evaluation of N_EV3_-adaptor for loading proteins onto circulating EVs *in vitro* and *in vivo*. (**a**) Schematic of N_GFP_-N_EV3_-mediated capture of GFP, loading onto EVs and EV-dependent cellular delivery *in vitro*. (**b**) Representative flow cytometry plots and quantification of GFP-positive EVs (n=3, Welch’s one-way ANOVA followed by Holm-adjusted pairwise Welch’s t-tests). N_GFP_-N_EV3_, GFP and EVs derived from 293T cells were mixed prior to flow cytometry. Blank refers to EVs alone. (**c**) Representative flow cytometry plots and quantification of GFP-positive cells (n=3, Welch’s one-way ANOVA followed by Holm-adjusted pairwise Welch’s t-tests). N_GFP_- N_EV3_-loaded EVs or a mixture of N_GFP_ and EVs were added to 293T cells together with GFP. GFP refers to cells treated with GFP alone; NC represents cells treated with PBS. (**d**) Schematic of the capture of circulating GFP by N_GFP_-N_EV3_-loaded EVs and subsequent delivery to the liver. (**e**) Representative *ex vivo* fluorescence images and quantification of GFP signals in body-wide tissues (n=4, two-sided unpaired Welch’s t-tests). GFP and N_GFP_-N_EV3_-loaded EVs were sequentially injected into *C57BL/6* mice intravenously and tissues were harvested 1 h after injection. H, heart; S, spleen; Lu, lung; Li, liver; K, kidney; NC refers to mice injected with buffer. (**f**) Schematic of protein A-N_EV3_-mediated loading of multiple antibodies onto EVs. (**g**) Representative flow cytometry plots and quantification of APC-Cy7- and FITC-positive EVs (n=3, two-sided unpaired Welch’s t-test). APC-Cy7-conjugated anti-CD45 antibody was incubated with WGA-AF488-labeled EVs derived from 293T cells in the presence or absence of protein A-N_EV3_. PA-N_EV3_ refers to protein A-N_EV3_ unless otherwise specified. NC refers to WGA-AF488-labeled EVs. Blank refers to unlabeled EVs. (**h**) Representative flow cytometry plots and quantification of FITC-, APC-Cy7-, APC-, BV421- and PE-Cy7-positive EVs (n=3, two-sided unpaired Welch’s t-test). Protein A- N_EV3_ was incubated with four fluorescent antibodies including APC-Cy7-anti-CD45, APC-anti-CD80, BV421-anti-XCR1 and PE-Cy7-anti-CD25, followed by co-incubation with WGA-AF488-labeled EVs derived from 293T cells. A mixture of the four antibodies and EVs was used as the negative control. (**i**) Quantification of FITC-IgG fluorescence in 293T cells and the supernatant after the addition of FITC-IgG, EVs and protein A-N_EV3_ or protein A-N_GFP_ (n=5, Welch’s one-way ANOVA followed by Holm-adjusted pairwise Welch’s t-tests). PA-N_GFP_ refers to protein A-N_GFP_ unless otherwise specified. (**j**) Circulating IgG levels in serum from *C57BL/6* mice injected with protein A-N_EV3_ (n=8), protein A-N_GFP_ (n=4) or saline (n=3) intravenously. Arrows indicate the injection times. Statistical significance was assessed by two-tailed t tests; *p < 0.05; **p< 0.01.

### Protein A-N_EV3_ improves survival and pathologies of SLE mice

Given the striking effect of protein A-N_EV3_ in reducing circulating IgG antibodies *in vivo*, we explored its therapeutic efficacy in SLE, a chronic, multisystem autoimmune disease characterized by elevated levels of autoantibodies such as double-stranded DNA (dsDNA) IgG^31^. We intravenously administered protein A-N_EV3_ at a dose of 2.5 mg/kg every three days for 10 weeks^32^ to *MRL/Lpr* mice (Fig. 6a), a mouse model resembling SLE phenotypes^33^. Circulating total IgG and anti-dsDNA IgG antibodies were significantly reduced in the blood of *MRL/Lpr* mice treated with protein A-N_EV3_ with a 100% survival rate, though only 25% of mice remained for protein A-N_GFP_ and PBS controls at 10 weeks post-treatment (Fig. 6b, c). Given that cachexia^34^, cervical lymphadenopathy^35^ and splenomegaly^36^ are common manifestations in SLE patients, measurement of these parameters revealed significant increase in body-weight (Fig. 6d), declines in cervical lymph node weight (Fig. 6e) and spleen size and weight (Fig. 6f) of mice treated with protein A-N_EV3_ compared with protein A-N_GFP_ and PBS control groups. Lupus nephritis (LN) is a severe complication caused by autoantibody complexes in SLE^37^, and examination of Urinary Albumin-to-Creatinine Ratio (UACR), a key proteinuria parameter reflecting SLE-associated LN^38^, showed a significant drop in urine from mice treated with protein A-N_EV3_ compared with protein A-N_GFP_ and PBS controls (Fig. 6g). Aligning with the UACR data, substantially reduced immune cell infiltration was observed in kidneys from mice treated with protein A-N_EV3_ compared to protein A-N_GFP_ and PBS controls, with no morphological changes found in liver (Fig. 6h). In summary, these findings demonstrate that protein A-N_EV3_ reduced circulating autoantibodies, prolonged survival and alleviated pathologies in SLE mice *in vivo*.

**Figure 6.**
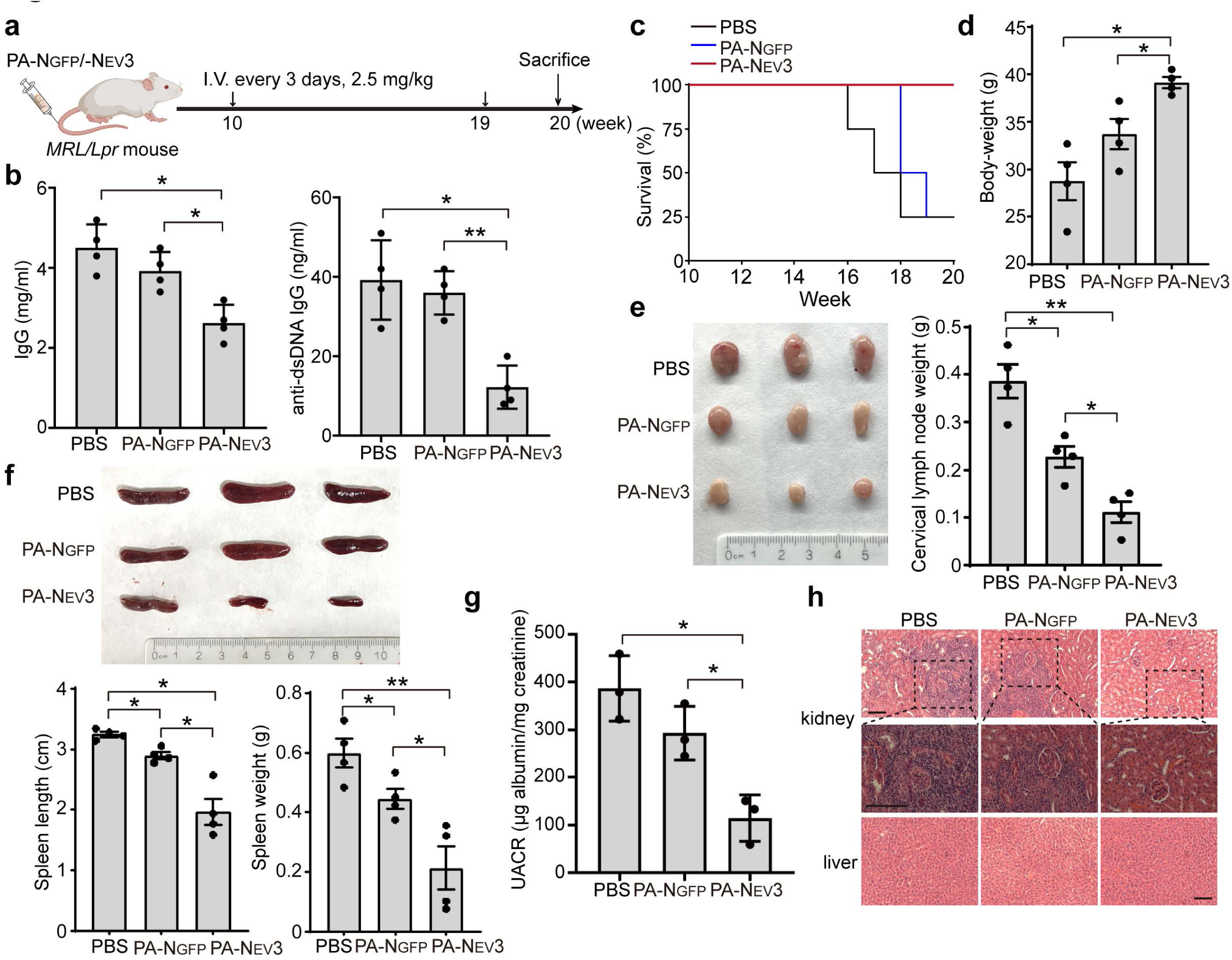
Evaluation of protein A-N_EV3_ in capturing circulating antibodies in SLE mice *in vivo*. (**a**) Diagram of the dosing regimen in *MRL/Lpr* mice. I.V. refers to intravenous injection. (**b**) Measurement of total IgG and anti-dsDNA IgG in serum from *MRL/Lpr* mice treated with protein A-N_GFP_ or protein A-N_EV3_ (n=4). PA-N_EV3_ or PA-N_GFP_ refers to protein A-N_EV3_ or protein A-N_GFP_ unless otherwise specified. (**c**) Kaplan-Meier survival curves for mice treated with PBS, protein A-N_GFP_ or protein A-N_EV3_. (**d**) Measurement of body-weight of *MRL/Lpr* mice treated with protein A-N_GFP_ or protein A-N_EV3_ at the experimental endpoint. (**e**) Representative images and weights of the largest cervical lymph nodes of treated mice (n=4). (**f**) Representative images, lengths and weights of spleens from treated mice at the experimental endpoint (n=4). (**g**) Measurement of urinary albumin-to-creatinine ratio (UACR) in treated mice (n=4). (**h**) Representative hematoxylin-and-eosin staining of kidney and liver sections (scale bar=100 μm). Statistical significance was assessed using Welch’s one-way ANOVA followed by Holm-adjusted pairwise Welch’s t-tests; *p < 0.05; **p< 0.01.

### Anti-TfR1-protein A-N_EV3_ redirects circulating EVs to muscle, resulting in phenotypic rescue of DMD mice

Inspired by the therapeutic outcomes in *SLE* mice, we wished to investigate protein A-N_EV3_ in muscle diseases as muscle-specific delivery of biotherapeutics remains one of the most significant challenges^39^. Considering that blood EVs were shown to improve the phenotypes in DMD mice by effectively blocking membrane rupture and restoring sarcolemmal localization of dystrophin-associated protein complex (DAPC), a large multicomponent complex disrupted and mislocalized as a result of membrane instability and leakage caused by loss of dystrophin in muscle cells^40^, we selected anti-TfR1 IgG, an antibody against TfR1 which is a receptor used for muscle-targeted delivery in DMD^21^, and formed anti-TfR1-protein A-N_EV3_ (Fig. 7a). After sequential intravenous injection of DiR-labeled EVs from mouse serum and anti-TfR1-protein A-N_EV3_ into wild-type *C57BL/6* mice, significantly higher fluorescence signals were seen in quadriceps, gastrocnemius and triceps and muscle sections from mice treated with anti-TfR1-protein A-N_EV3_ compared with anti-TfR1-protein A-N_GFP_, though no significant difference was observed in liver and kidney (Fig. 7b, c), confirming anti-TfR1-protein A-N_EV3_’s ability to redirect circulating EVs to muscle. Significantly greater muscle strength (Fig. 7d) and lower levels of serum creatine kinase (Fig. 7e), an enzyme markedly elevated in DMD patients due to chronic muscle damage^41^, were detected in *mdx* mice treated with intravenous administration of anti-TfR1-protein A-N_EV3_ at a dose of 10 mg/kg every three days for a week than anti-TfR1-protein A-N_GFP_ under identical conditions^32^. Restoration of DAPC was detected on the sarcolemma of quadriceps from *mdx* mice treated with anti-TfR1-protein A-N_EV3_ compared with anti-TfR1-protein A-N_GFP_ (Fig. 7f, g). Sarcolemma integrity was dramatically improved as demonstrated by reduced levels of IgG (Fig. 7h), a marker of impaired sarcolemmal integrity^40^, and fibrosis (Fig. 7i) in quadriceps of *mdx* mice treated with anti-TfR1-protein A-N_EV3_ compared with anti-TfR1-protein A-N_GFP_. Taken together, these data indicate that anti-TfR1-protein A-N_EV3_ is able to hijack and redirect circulating EVs to muscle, resulting in improved phenotypes of DMD mice *in vivo*.

**Figure 7.**
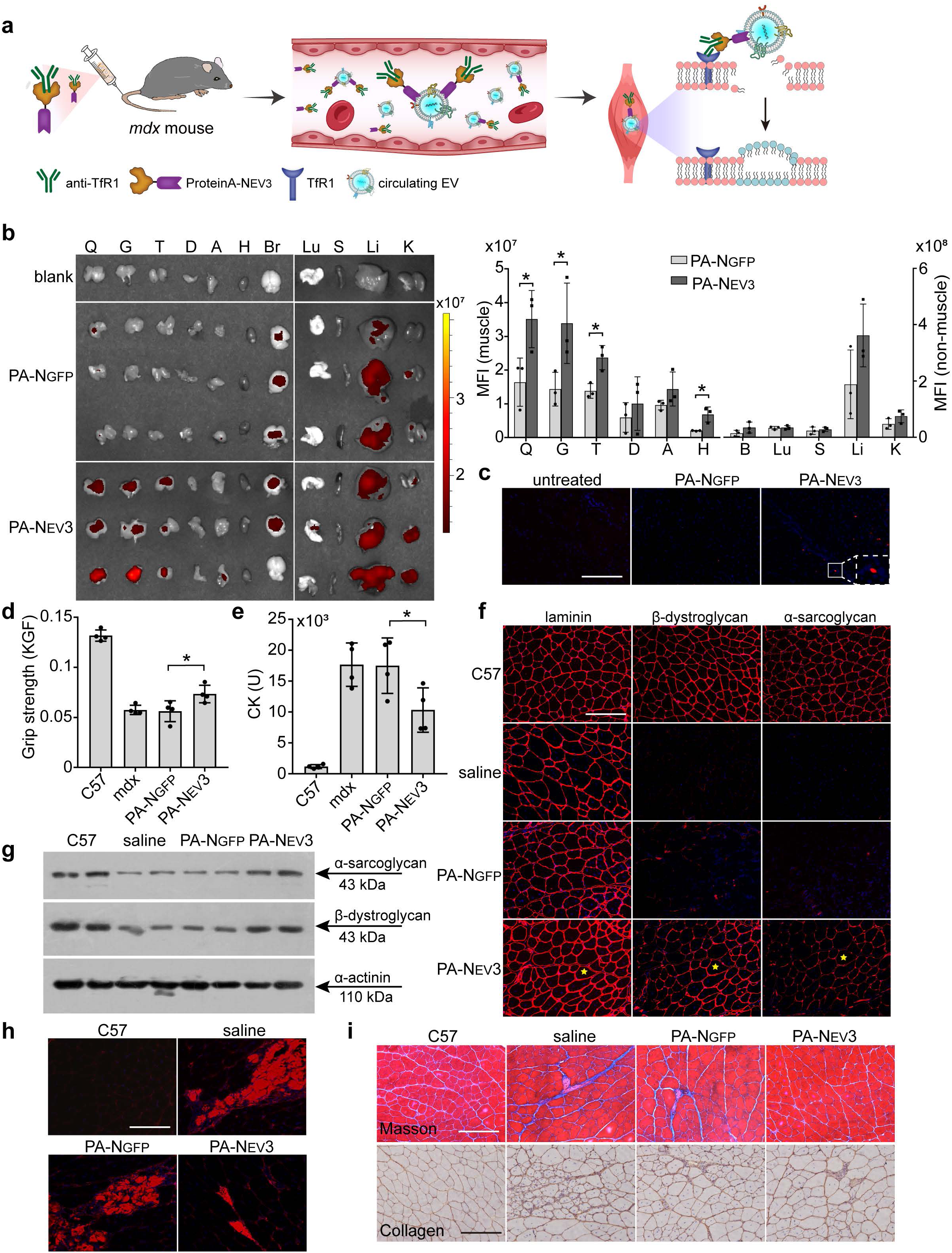
*In vivo* assessment of anti-TfR1-protein A-N_EV3_ in redirecting endogenous EVs to skeletal muscle in *mdx* mice. (**a**) Schematic of anti-TfR1-protein A-N_EV3_-mediated recruitment of circulating EVs to skeletal muscle. (**b**) Representative *ex vivo* fluorescence images and quantification of DiR signals in body-wide tissues (n=3, two-sided unpaired Welch’s t-tests). DiR-labeled EVs and anti-TfR1-protein A-NEV3 or anti-TfR1-protein A-NGFP were sequentially injected into *C57BL/6* mice intravenously. Q, quadriceps; G, gastrocnemius; T, triceps; D, diaphragm; A, abdominal muscle; H, heart; Br, brain; Lu, lung; S, spleen; Li, liver; K, kidney. (**c**) Representative fluorescence microscopy images of quadriceps sections from treated mice (scale bar=100 μm). Red dots are DiR signals. Measurement of grip strength (**d**) and levels of serum creatine kinase (CK) (**e**) in *mdx* mice treated with anti-TfR1-protein A-N_EV3_, anti-TfR1-protein A-N_GFP_ or untreated controls (n=4, two-sided unpaired Welch’s t-tests with Holm correction). Anti-TfR1-protein A-N_EV3_ or anti-TfR1-protein A-N_GFP_ was intravenously injected into *mdx* mice at the dose of 10 mg/kg every 3 days for 1 week. (**f**) Representative immunofluorescence images of laminin, β-dystroglycan and α-sarcoglycan in quadriceps from treated *mdx* mice (scale bar=100 μm). (**g**) Western blot analysis of α-sarcoglycan and β-dystroglycan in quadriceps from treated *mdx* mice. α-actinin was used as the loading control. A total of 50 μg protein was loaded per lane. (**h**) Representative immunofluorescence images of IgG distribution in quadriceps from treated *mdx* mice (scale bar=100 μm). (**i**) Representative Masson’s trichrome and immunohistochemical staining of collagen in quadriceps from treated *mdx* mice (scale bar=100 μm). Statistical significance was assessed by two-tailed t tests; *p < 0.05.

## Discussion

Here, we demonstrated that N_EV3_, as an EV-anchoring nanobody and a modular adaptor, enables efficient loading of a wide spectrum of protein therapeutics onto EVs via fibronectin and manipulation of circulating EVs after systemic administration *in vivo*. The central advance is not simply an additional EV-binding ligand; rather, this strategy transforms the therapeutic protein construct from *ex vivo* EV manufacture to N_EV3_-containing proteins by engaging EVs already present in the circulation and combining them with independent cargo-binding or tissue-targeting modules. Across reporter, RNA-binding and genome-editing cargoes, IgG recruitment and two disease models (Fig.8), the data establish the feasibility of this *in situ* loading and redirection concept with an EV-anchoring nanobody. For SLE, the intrinsic liver tropism of circulating EVs was harnessed as delivery vehicles to reduce levels of total and anti-dsDNA IgG antibodies in circulation by simply administering protein A-N_EV3_. In the case of DMD, a “killing two birds with one stone” effect was achieved by exploiting N_EV3_’s EV-anchoring and anti-TfR1’s muscle-targeting abilities, thus accomplishing muscle-specific delivery of circulating EVs as therapeutic agents via intravenous injection of anti-TfR1-protein A-N_EV3_ in *mdx* mice. To our knowledge, this is the first demonstration showcasing the ability of an EV-anchoring nanobody in maneuvering endogenous circulating EVs for different therapeutic purposes.

**Figure 8.**
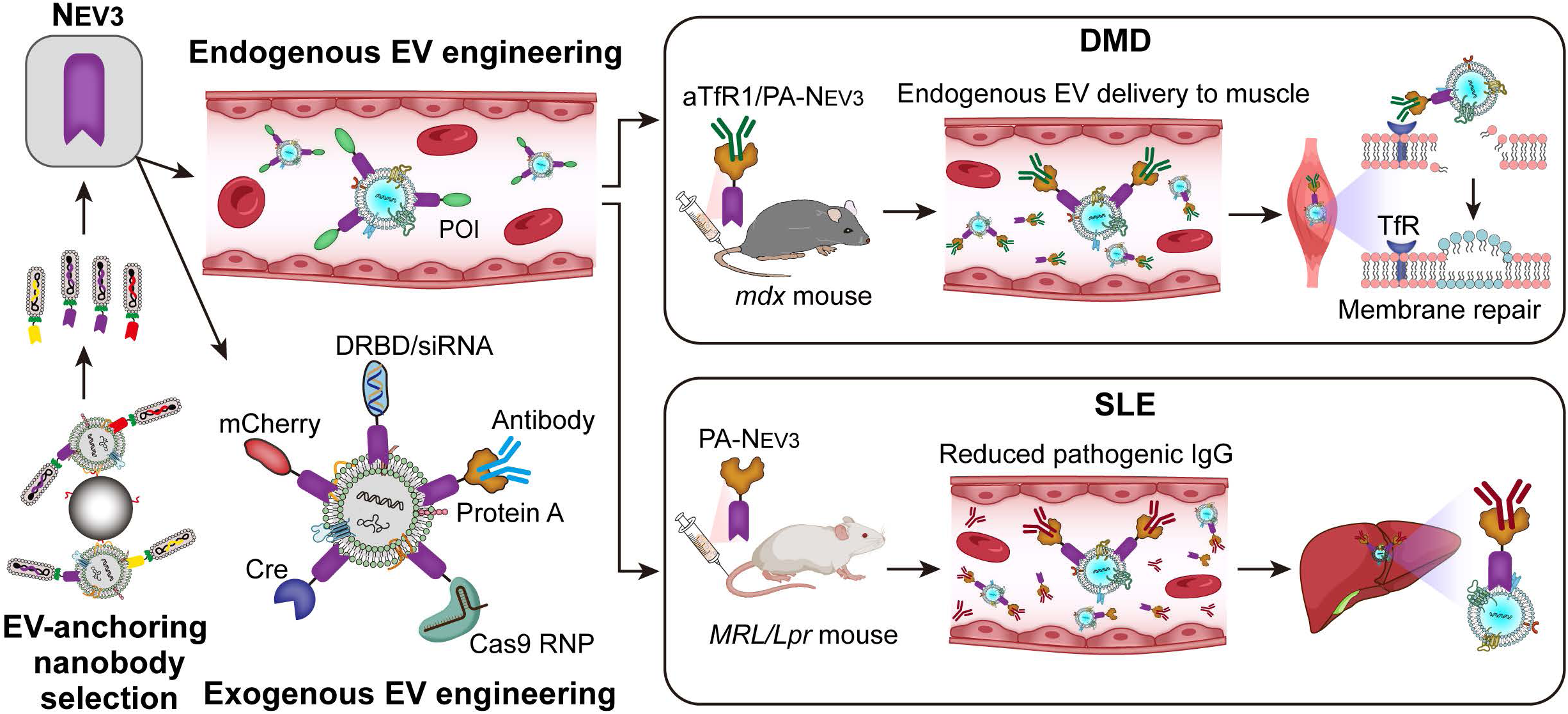
Schematic for N_EV3_-mediated loading of different therapeutic cargoes onto exogenous or endogenous EVs and simultaneous capture of circulating protein and EVs *in vitro* and *in vivo*. POI refers to Protein Of Interest.

The breadth of the cargo experiments suggests that N_EV3_ can accommodate proteins with distinct sizes and functions, including DRBD, Cre and Cas9. These highly encouraging data proved the potency of N_EV3_ in effectively loading different protein therapeutics on EVs and can be further exploited in gene therapy with minimal immunogenicity concerns, particularly when EVs from patients own are employed. Although detailed studies on cargo stoichiometry across EV subpopulations, cytosolic escape efficiency, repeated dose pharmacology or immunogenicity are warranted prior to clinical deployment.

The superior binding affinity of protein A for IgG antibodies^42^ makes it a convenient proof-of-concept IgG-binding module, but its bacterial origin and broad Fc affinity^43^ make it unsuitable as the present therapeutic format. Mammalian IgG binders such as Fc γ receptors (FcγRs)^44^ or engineered human IgG-binding domains may offer more translatable alternatives, although their selectivity, Fc-receptor interactions and immunological consequences will require direct testing. In the SLE study, the reduction in autoantibodies and disease measures is encouraging, though more extensive studies with the fate of recruited IgG, liver uptake and comprehensive mechanistic elucidation are warranted in future with a mammalian version of IgG binder.

Endogenous circulating EVs were shown to mitigate symptoms in different pathological conditions including blocking leaky muscle membrane in DMD^40^, promoting angiogenesis and improving heart function in infarcted heart tissues^45^, and enhancing fast vascularization in critical mandibular defects^46^ or wound healing^47^ *in vivo* by localized or passive delivery. Here, the muscle-targeting experiment illustrates a different scenario by navigating endogenous EVs to muscle via intravenous injection of anti-TfR1-protein A-N_EV3_, thus accomplishing tissue-specific accumulation of endogenous circulating EVs with resultant therapeutic effects in DMD mice. This delivery strategy can be broadly applied to different tissue-targeting moieties or any moiety of interest for different therapeutic applications, thus transforming EV-based to protein-based therapeutics.

Although fibronectin is often defined as a constituent of the extracellular matrix that is involved in different biological processes such as cell adhesion and differentiation in both soluble and cell-associated forms^48^, it also has been extensively found on the surface of EVs from different sources including hepatocytes, melanocyte, fibroblasts and tumor cells^25,26,49,50^, indicating the abundance of fibronectin on EVs of different sources. Also, elegant verification of direct association of fibronectin with EVs was established by immunoprecipitation, proteinase protection and fibronectin ablation^49^ as we did in this study. We measured the binding affinity of N_EV3_ with intact EVs based on the consideration that the purified fibronectin components might differ from EV-associated forms, nevertheless, more extensive studies including purified components and mutational validation of the predicted interface are needed to define the binding specificity.

In summary, our study demonstrates that N_EV3_ enables modular binding of protein constructs with EVs via fibronectin and *in situ* loading of biologics on and tissue redirection of circulating vesicles *in vivo*, thus providing a tool for protein engineering of EVs *in vitro* and *in vivo* and targeted delivery of protein therapeutics. N_EV3_ allows hijacking of endogenous circulating EVs either as delivery vehicles or as therapeutic agents in SLE and DMD models as proof-of-concept, showcasing endogenous EVs as a recruitable pharmacological compartment.

## Methods

### Ethics statement

Animal experiments were conducted at the Animal Unit of Tianjin Medical University (Tianjin, China) according to procedures approved by the institutional ethics committee (Permit Number: SYXK2023-0004). All procedures were conducted in accordance with institutional guidelines for the care and use of laboratory animals.

### Animals and injections

Eight-week-old female *MRL/lpr* mice were purchased from SPF Biotechnology Co., Ltd. (Beijing, China). Six-week-old male *mdx* mice were purchased from The Jackson Laboratory (US) and maintained in the laboratory. Cre reporter mice (mTmG) were kindly provided by Prof. Xiongwen Chen (Tianjin Medical University). Age- and sex-matched *C57BL/6* mice were purchased from Beijing Vital River Laboratory Animal Technology Co., Ltd. (Beijing, China) and used as wild-type controls wherever appropriate. The number of mice used for each experiment is specified in the corresponding figure legends.

For local intramuscular injection, Cre-N_EV3_ (5 μg) was incubated with EVs derived from 293T cells (10 μg) and injected into the right tibialis anterior (TA) muscles of loxP-tdTomato-stop-loxP-GFP mice. As a control, Cre-N_GFP_ incubated with EVs derived from 293T cells was injected into the left TA muscle of the same mice. TA muscle tissues were collected at 1 week post-injection and imaged using an IVIS Spectrum imaging system (PerkinElmer, US).

For the systemic lupus erythematosus experiment, protein A-N_EV3_ or protein A-N_GFP_ was administered intravenously to 10-week-old female *MRL/lpr* mice at a dose of 2.5 mg/kg every three days for 10 weeks and tissues were harvested 1 week after the last injection. For the DMD therapeutic experiment, protein A-N_EV3_ or protein A-N_GFP_ was mixed with anti-transferrin receptor (anti-TfR) antibodies at a 1:1 molar ratio and incubated for 2 h at 4°C. The resulting complex was intravenously administered to 6-week-old *mdx* mice at a dose of 10 mg/kg every 3 days for 1 week and tissues were harvested 2 weeks after the last injection. Mice were euthanized by CO₂ inhalation or cervical dislocation at the designated timepoints and tissues were either snap-frozen in liquid nitrogen-cooled isopentane and stored at -80°C or fixed with Bouin’s solution (Sigma, St. Louis, MO, US) and embedded in paraffin for immunohistochemical and histological studies.

For the inhibition of EV release *in vivo*, GW4869 (MCE, US) was intraperitoneally injected into *C57BL/6* mice at a dose of 2.5 mg/kg once daily for five consecutive days and the buffer containing 0.001% DMSO was used as control. Blood was immediately collected after the final administration for subsequent analysis of circulating EVs.

### Cell culture

Murine C2C12 cells (mouse myoblasts), Hepa1-6 and dendritic cells (DC2.4), human embryonic kidney 293T (293T) and pancreatic cancer cells (Panc02) were cultured as previously reported^51^. Briefly, cells were grown at 37°C in 5% CO_2_ in Dulbecco’s modified Eagle’s medium (DMEM) supplemented with 10% fetal bovine serum (FBS) and 1% penicillin and streptomycin. Myotubes were obtained from confluent C2C12 cells seeded in gelatin-coated 12-well plates after 3 days of serum deprivation at 37°C under a 5% CO_2_ atmosphere in DMEM with 2% horse serum (Hyclone, US). Mouse Hepa1-6 cells were purchased from Boster Biological Technology, Ltd (Wuhan, China) and cultured in DMEM medium with 2 mM glutamine and 10% FBS as per manufacturer’s instructions. DCs were cultured in DMEM supplemented with 10% FBS, 1% PS, 1% glutamine and 50 μM β-mercaptoethanol. FBS was centrifuged at 100,000 × g for 70 min to remove EVs present in FBS prior to addition to cells for EV production. Cell supernatants of human mesenchymal cells were kindly provided by SHUNHO CELL BIOLOGY TECHNOLOGY Corp. (Tianjin, China).

### Generation of *fibronectin* knockout and reporter cells

To generate *fibronectin* knockout 293T cells, we cloned the *fibronectin* sgRNA (sgFN1) (5’-GCAAGCGTGAGTACTGACCG-3’) into the Lenti-CRISPRv2 plasmid (Addgene Inc., US), with the resulting LentiCRISPRv2-sgFN1-puro containing hSpCas9 and the chimeric guide RNA. To obtain lentiviruses expressing Cas9 RNP, 293T cells were seeded in a 10-cm Petri dish at a density of 5× 10^7^ cells and cultured for 24 h, followed by co-transfection with LentiCRISPRv2-*fibronectin* sgRNA-puro, ΔR and VSVG plasmids at a mass ratio of 2:1:1 (µg) by calcium phosphate precipitation. Viruses were harvested and titred 48 h later and used for subsequent infection of 293T cells. 293T cells were seeded in a 12-well plate and infected with 5× 10^7^ PFU of lentivirus-expressing Cas9 RNP for 12 h. The infection was repeated three times at 24 h intervals, followed by puromycin selection (1 µg/ ml) for 2 weeks. To generate Cre reporter 293T cells, the Cre reporter plasmid (CMV-mCherry-LoxP-stop-LoxP-GFP-puro) was synthesized by Genewiz (Suzhou, China), followed by production of lentiviruses containing the expression cassettes, and infection and selection of 293T cells with puromycin (1 µg/ ml) for 2 weeks as described above. mCherry-positive cells were sorted by fluorescence-activated cell sorting (FACS) (BD FACSAria Fusion, New Jersey, US) for subsequent tests. To generate Cas9 reporter 293T cells, the CRISPR Cas9 reporter plasmid purchased from Beijing Zhongyuan Ltd. (Beijing, China) was used to generate lentiviruses, followed by infection and selection of 293T cells as described above.

### *In vitro* biopanning

Our proprietary synthetic nanobody library with humanized framework region backbones was used for *in vitro* biopanning experiments. EVs derived from differentiated mouse myotubes were immobilized on CP05-coated magnetic beads (provided by Dr Yiqi Seow, Genome Institute of Singapore, A*STAR, Singapore). Briefly, 50 μg EVs from differentiated mouse myotubes were incubated with 20 μl CP05-coated magnetic beads at 4°C for 30 min. Phages (1×10^12^ pfu, plaque-forming units) were incubated with EV-bound beads at 4°C for 2 h during rounds 1 to 3, and reduced to 1 h for rounds 4 to 7. Unbound phages were removed by washing 12 times with phosphate buffered saline (PBS) containing 0.1% Tween-20 briefly. Bound phages were eluted using 0.1 M glycine-HCl (pH 2.2), and the eluate was immediately neutralized with 1 M Tris-HCl (pH 8.8)^18^. Eluted phages were amplified using VCSM13 helper phage and used as the input for the subsequent round of selection. After seven rounds of selection, twenty phage clones were randomly selected and sequenced by Genewiz (Beijing, China).

### Preparation and purification of EVs

Cell culture medium was sequentially centrifuged at 2, 000 × g for 30 min, followed by 10,000 × g for 30 min to remove cells and cellular debris. The supernatant was collected and filtered with a 0.22 µm filter (Millex, Germany), followed by ultracentrifugation at 100,000 × g for 70 min to pellet EVs. The EV pellets were washed in a large volume of PBS and recovered by centrifugation at 100,000 × g for 70 min. The total protein concentration was quantified by the Bradford assay (Sangon Biotech, China). For labeling, purified EVs were incubated with 1 μM DiR (1,1′-dioctadecyl-3,3,3′,3′-tetra methy =lindotricarbocyanine iodide; Invitrogen, US) or WGA-Alexa Fluor 488 (WGA-AF488; MedChemExpress LLC, US) at 37°C for 45 min. Labeled EVs were subsequently ultracentrifuged at 100,000 × g for 70 min to remove unbound dye^52^. The resulting EV pellets were resuspended in PBS for subsequent experiments.

### Characterization of EVs

N_EV3_ (2 μg) was incubated with EVs (10 μg) derived from differentiated mouse myotubes at 4°C for 1 h to obtain N_EV3_-modified EVs (EVN_EV3_). The size distribution of EVs and EVN_EV3_ was measured by nanoparticle tracking analysis (NTA) (NanoSight, Malvern, UK) as described previously^18^. For all recordings, we used a camera level of 13-14 and automatic functions for all post-acquisition settings except for the detection threshold, which was fixed at 5. All samples were diluted in 0.22µm filtered PBS at ratios of 1:100 and 1:1000 to achieve a particle concentration of between 1 × 10⁸ and 1 × 10⁹ particles/ml. The camera focus was adjusted to make the particles appear as sharp dots. Using the script control function, three 60s videos for each sample were recorded, incorporating a sample advance and a 5s delay between each recording. Morphology was visualized using a high-resolution transmission electron microscope (TEM, Hitachi HT7700, Tokyo, Japan). Briefly, the resuspended EVs were diluted in water (1 μg/μl) and mixed with an equal volume of 4% paraformaldehyde (PFA). EVs were adsorbed onto a glow-discharged, carbon-coated formvar film attached to a metal specimen grid. Excess solution was blotted off and the grid was immersed with a small drop (50 μl) of 1% of glutaraldehyde for 5 min followed by washing with 100 μl distilled water for 8 times (2 min each time). Subsequently, the grid was transferred to 50 μl uranyl-oxalate solution (pH 7.0) for 5 min and then to 50 μl methyl cellulose-uranyl acetate (100 μl 4% uranyl acetate and 900 μl 2% methyl cellulose) for 10 min on ice. The excess solution was blotted off and the sample was dried and examined by TEM^18^.

### Recombinant protein purification and labeling

The cDNA sequences of N_EV3_, N_GFP_ and protein A-N_EV3_ were cloned into the pGEX-6P-1 plasmid at the downstream of a C-terminal GST tag (Supplementary Table S1), respectively. The mCherry, N_EV3_-mCherry, mCherry-N_EV3_, mCherry-N_GFP_, GFP, Cre-N_EV3_, Cre-N_GFP_ and DRBD-N_EV3_ sequences were cloned into the pET-28a vector at the downstream of a C-terminal His tag. The Cas9 and Cas9-N_EV3_ sequences were cloned into the pET-28b vector at the downstream of a C-terminal His tag (Supplementary Table S1). These plasmids were transformed into *E. coli* BL21 (DE3) (Zomanbio, China). All recombinant proteins were expressed in *E. coli* BL21 (DE3) and bacteria were grown in Luria-Bertani (LB) medium supplemented with kanamycin or ampicillin. Cultures were incubated at 37°C with shaking at 200 rpm until the optical density at 600 nm reached 0.6-0.8. Protein expression was then induced by adding isopropyl β-d-1-thiogalactopyranoside (IPTG; Solarbio, China) at a final concentration of 0.5 mM, followed by incubation at 16°C with shaking at 150 rpm for 16 h. The recombinant proteins with a GST tag were purified with Glutathione Agarose resin (Cytiva, US). The bound proteins were eluted with reduced glutathione (GSH, Solarbio, China) to retain the GST tag or cleaved using HRV3C protease to remove the GST tag. The recombinant proteins with His tags were incubated with His-tag magnetic beads (Beaver, China), and the bound proteins were eluted using imidazole (Solarbio, China). Before elution, the Cas9 and Cas9-N_EV3_ bound proteins were washed three times for 5 min each time with a high-salt buffer (25 mM Tris-HCl, 2 M NaCl, pH 8.0). Proteins were further purified using an AKTA FPLC system (Cytiva, US) equipped with an ion-exchange column (cation or anion exchange depending on protein properties) or a heparin column (used to purify DRBD-N_EV3_ to eliminate residual host nucleic acid impurities). Elution was performed using a salt gradient adjusted for each protein. SDS-PAGE was used to verify the purity of purified proteins after Coomassie Brilliant Blue staining.

For the labeling, proteins were first buffer-exchanged into 0.1 M sodium carbonate-sodium bicarbonate buffer (pH 9.0), and the final protein concentration was adjusted to 1 μg/μl. FITC (Fluorescein-5-isothiocyanate; Sigma-Aldrich, Germany) or Cy5 (Med-ChemExpress, US) dye was added to the protein solution at a dye-to-protein mass ratio of 1:10, and the mixture was thoroughly vortexed. The labeling reaction was carried out at 4°C overnight with protection from light. Unbound dyes were removed by dialysis against the buffer used for the corresponding dye.

### N_EV3_ binding assay *in vitro*

For the direct EV binding, 2 μg N_EV3_ or N_EV3_ fusion protein was incubated with 5 μg EVs derived from different sources (as specified in the corresponding figure legend) in 100 μl PBS at 4°C for 1 h protected from light, unless otherwise specified. Unbound protein was removed using a 100-kDa molecular-weight-cutoff centrifugal filter (Millipore, US), and the resulting EV-protein complexes were analyzed by flow cytometry as described previously^18^. For the comparison with mCherry placed either at the N- or C-terminus of N_EV3_, 5 μg EVs derived from 293T cells were incubated with 0.5, 1 or 2 μg N_EV3_-mCherry or mCherry-N_EV3_ under the same condition as described above. For measuring the binding of N_EV3_ to fibronectin-depleted EVs, mCherry N_EV3_ (2 μg) was incubated with WGA-AF488-labeled EVs derived from 293T cells (5 μg) or from *fibronectin*-knockout 293T cells (5 μg) in 100 μl PBS at 4°C for 1 h protected from light. Then, the binding was analyzed by flow cytometry.

For adaptor-mediated cargo loading experiments, FITC-labeled siRNA (0.25 μg) was incubated with DRBD-N_EV3_ (5 μg) on ice for 30 min, followed by co-incubation with 20 μg EVs derived from 293T cells in 100 μl PBS at 4°C for 1 h; 5 μg GFP was incubated with 5 μg N_GFP_-N_EV3_ on ice for 30 min, followed by co-incubation with 20 μg EVs derived from 293T cells in 100 μl PBS at 4°C for 1 h; 0.2 μl APC-Cy7-conjugated anti-CD45 antibody (Biolegend, US) was incubated with 1 μg protein A- N_EV3_ on ice for 30 min, followed by co-incubation with 5 μg WGA-AF488-labeled EVs derived from 293T cells in 100 μl PBS at 4°C for 1 h. For simultaneous loading, 0.05 μl APC-Cy7-conjugated anti-CD45, APC-conjugated anti-CD80, BV421-conjugated anti-XCR1 and PE-Cy7-conjugated anti-CD25 (Biolegend, US) were mixed with 1 μg protein A-N_EV3_ on ice for 30 min, followed by co-incubation with 5 μg WGA-AF488-labeled EVs derived from 293T cells in 100 μl PBS at 4°C for 1 h. The assembled complexes were analyzed by flow cytometry (BD FACS Verse, BD, US).

### N_EV3_ binding assay *in vivo*

To assess the binding of N_EV3_ to circulating EVs, 300 μg mCherry-N_EV3_ or mCherry- N_GFP_ was intravenously injected into *C57BL/6* mice. Blood was collected into EDTA-containing tubes 2 h post-injection and centrifuged at 1,000 × g for 20 min at 4°C to separate the plasma. Plasma EVs were isolated by ultracentrifugation and washed with PBS as described above. The mCherry fluorescence signal associated with isolated EVs was subsequently detected by flow cytometry. For the *in vivo* half-life assay, 300 μg mCherry-N_EV3_ or mCherry-N_GFP_ was injected into *C57BL/6* mice via the tail vein. Blood was collected at various time points (0, 1, 2, 4, 6 and 8 h), and plasma was recovered from the samples. For the fluorescence imaging, 50 μl plasma was mixed with 50 μl PBS and added to a black 96-well plate, and images were acquired using an IVIS Spectrum imaging system (PerkinElmer, US). Separately, 20 μl plasma was mixed with 80 μl PBS and added to a 96-well plate for fluorescence intensity measurement using a microplate reader (Revvity, US). The plasma half-life of mCherry-N_EV3_ was determined using a one-phase decay model in GraphPad Prism (GraphPad Software, Inc., US).

### Gel migration assay

DRBD-N_EV3_ (1, 5, 10 or 20 μg) was mixed with 1 μg *GAPDH* siRNA (UGACCUCA ACUACAUGGUUTT)^28^. The reaction mixtures were adjusted to a final volume of 20 μl with PBS and incubated on ice for 30 min. Following incubation, the samples were subjected to gel electrophoresis at 180 V for 8 min. The gels were then visualized using a gel documentation system (Tanon 1600, China).

### Cas9-RNP assembly *in vitro*

A target sgRNA sequence of GGGCCACTAGGGACAGGAT was generated via *in vitro* transcription using the HiScribe™ T7 Quick High Yield RNA Synthesis Kit (NEB, US) according to the manufacturer’s instructions. To form the secondary scaffold structure, 500 ng sgRNA was subjected to a gradient annealing program (75°C for 5 min, followed by 0°C for 10 min). The sgRNA was then incubated with 2 μg Cas9- N_EV3_ or Cas9 in the RNP assembly buffer (25 mM Tris-HCl, 200 mM KCl, 2 mM DTT, 5 mM MgCl_2_, 5% glycerol) at 37°C for 15 min.

### Fluorescence microscopy

For the Cre-N_EV3_ experiment, Cre reporter 293T cells were seeded in 24-well plates containing sterile coverslips at a density of 1× 10^5^ cells per well and cultured overnight. Cre-N_EV3_ or Cre-N_GFP_ (10 μg) was incubated with EVs derived from 293T cells (20 μg) and then added to Cre reporter 293T cells. Cells were observed at 72 h post-incubation using a confocal fluorescence microscope (Olympus, FV1000, Japan). For the Cas9-N_EV3_ experiment, Cas9 reporter cells were seeded in 24-well plates containing sterile coverslips at a density of 1× 10^5^ cells per well and cultured overnight. Subsequently, the assembled Cas9-RNP was incubated with 20 μg EVs derived from 293T cells at 4°C for 2 h. The RNP-EV complexes were then added to Cas9 reporter cells and fluorescence signals were visualized at 48 h post-incubation using a confocal fluorescence microscope.

### Flow cytometry

293T cells were seeded in 24-well plates at a density of 2× 10^5^ cells per well and cultured overnight. To evaluate the ability of N_GFP_-N_EV3_ to capture GFP in the supernatant, N_GFP_-N_EV3_ (10 μg) was incubated with EVs derived from 293T cells (20 μg) at 4°C for 2 h. Then, GFP (10 μg) was added to 293T cells seeded in culture plates and mixed well, followed by the addition of the mixture of N_GFP_-N_EV3_ and EVs. Cells were further incubated for 24 h and washed with cold PBS briefly and digested with trypsin, followed by washing and resuspension in cold PBS, prior to analysis using a BD Verse flow cytometer (BD Biosciences, US). To measure cellular uptake, mCherry- N_EV3_, mCherry-N_EV3_ or mCherry-N_GFP_ (5 μg) was incubated with WGA-AF488-labeled EVs derived from 293T cells (10 μg) at 4°C for 2 h, followed by the addition of the resulting mixture to 293T cells and incubation for 24 h before flow cytometry (BD FACS LSRFortessa, BD, New Jersey, US). For siRNA-DRBD-N_EV3_, FITC-labeled siRNA (1 μg) was incubated with DRBD-N_EV3_ (10 μg) on ice for 30 min, followed by co-incubation with EVs derived from 293T cells (20 μg), or siRNA alone with EVs at 4°C for 2 h. The mixture of siRNA or siRNA-DRBD-N_EV3_ with EVs was added to 293T cells and incubated for 24 h before flow cytometric analysis. For the Cas9 experiment, Cy5-labeled Cas9-N_EV3_ or Cy5-labeled Cas9 (5 μg) was incubated with EVs derived from 293T cells (20 μg) at 4°C for 2 h. The mixture was added to 293T cells and co-incubated for 24 h before flow cytometric analysis.

For *in vivo* evaluation of endogenous EVs, 300 μg mCherry-N_EV3_ or mCherry-N_GFP_ protein was administered intravenously to *C57BL/6* mice. Liver tissues were harvested and dissociated using collagenase after perfusion 2 h post-injection as described previously^18^. The digested liver tissue was passed through a 70-μm cell strainer to collect the cell suspension. The cell suspension was then centrifuged at 50 × g for 5 min, and the resulting pellet was collected as hepatocytes. Subsequently, the remaining supernatant was centrifuged at 4,500 rpm for 5 min to collect other cell pellets for antibody staining. The pelleted cells were subsequently stained with APC-Cy7-anti-mouse CD45 (1:500), PE-anti-mouse CD31 (1:200), BV421-anti-mouse CD11b (1: 200), and FITC-anti-mouse F4/80 (1:200; BioLegend, US). Endothelial cells were identified as CD45⁺CD31⁺ and macrophages were gated as CD45⁺CD11b⁺F4/80⁺ cells.

### Enzyme-Linked Immunosorbent Assay (ELISA)

For IgG binding *in vitro*, 293T or Hepa1-6 cells were seeded in 96-well plates at a density of 1 ×10^4^ cells per well and cultured for 24 h. The culture medium was then replaced with EV-depleted medium. FITC-labeled rabbit anti-mouse IgG (1:1,000; Thermo Fisher Scientific, US), protein A-N_EV3_ or the control protein A-N_GFP_ (1 μg per well), and EVs derived from 293T cells (2.5 μg per well) were added separately to the culture medium. After incubation for 24 h, the culture supernatants were collected and cells were washed 5 times with DPBS briefly and lysed in RIPA lysis buffer (Sangon Biotech, China). FITC fluorescence in the culture supernatants and cell lysates was measured using a microplate reader (PerkinElmer, US).

### Super-resolution microscopy

Super-resolution microscopy (ONI, Oxford, UK) was performed to measure the number of N_EV3_ molecules associated with single EVs as per manufacturer’s instructions. Briefly, 2.5 µl EVs derived from 293T cells (1×10⁹ particles/µl) were blocked with 3.5 µl blocking solution for 5 min, followed by staining with 1 µl anti-CD9-561, anti-CD63-561 and anti-CD81-561 (provided in the kit) or 1 μg Cy5-labeled N_EV3_ overnight at 4°C. The stained EVs were added to the pre-treated lanes on the plate, followed by washing with the washing buffer to remove free antibodies or N_EV3_. The lanes were imaged with a Nanoimager (ONI, Oxford, UK), and the image processing and data analysis were performed using the EV Profiling program on an online platform (https://alto.codi.bio/).

### AuNP synthesis and blocking

Gold nanoparticles (AuNPs) were synthesized using a citrate reduction method^53^. Briefly, 50 ml 1 mM chloroauric acid solution (Solarbio, China) was heated to boiling under vigorous stirring. Subsequently, 5 ml 38.8 mM trisodium citrate solution (Solarbio, China) was rapidly added, and the reaction mixture was continuously boiled and stirred for 15-20 min until the color changed from pale yellow to wine red. The reaction mixture was then cooled to room temperature (RT) under continuous stirring and stored at 4°C in the dark. AuNPs were incubated with DiR-labeled EVs derived from differentiated C2C12 cells at EV-to-Au mass ratios of 1:10, 1:20 and 1:40 at RT for 30 min. BSA (1μg/μl) was added and incubated at RT for 30 min to block the unoccupied AuNP surface. Subsequently, 2 μg mCherry-N_EV3_ was co-incubated with 10 μg AuNP-coated EVs in 100 μl PBS at 4°C for 2 h. Binding was subsequently quantified by flow cytometry.

### Enzyme digestion

For proteinase K digestion, 1 μg/μl proteinase K (Sigma-Aldrich, Germany) was added to 1 μg/μl WGA-labeled EVs derived from differentiated C2C12 cells. The mixture was gently mixed and incubated at 37°C for 30 min, and the digestion was terminated by 1 mM protease inhibitor phenylmethylsulfonyl fluoride (PMSF; Solarbio, China). Then, the proteinase K digested EVs or untreated EVs (20 μg) were co-incubated with mCherry-N_EV3_ (2 μg) in 100 μl PBS at 4°C for 1 h. The binding efficiency was subsequently quantified by flow cytometry. For proteinase K and PNGase F glycosidase digestion, 1 μg/μl DiR-labeled EVs derived from differentiated C2C12 cells were incubated with 0.5 U/ml PNGase F glycosidase (Sigma-Aldrich, Germany) at 37°C for 16 h, followed by the addition of 1 μg/μl proteinase K and further incubation at 37°C for 30 min. Subsequently, 1 mM PMSF was added to terminate the digestion. Digested EVs or untreated EVs (20 μg) were co-incubated with of mCherry-N_EV3_ (2 μg) in 100 μl PBS at 4°C for 1 h. The binding efficiency was subsequently quantified by flow cytometry.

### Immunoprecipitation

For the immunoprecipitation of EV lysates, 50 μg GST-N_EV3_ protein and 200 μg EVs derived from differentiated C2C12 cells were co-incubated in 200 μl PBS at 4°C for 2 h. Subsequently, 0.1% Triton X-100 was added to lyse EVs for 1 h, followed by the addition of 20 μl Glutathione Agarose resin (GST-beads) and incubation overnight at 4°C. As negative controls, EV lysate or GST-_NEV3_ protein alone was incubated with GST-beads. Next day, 1 ml PBS was added and the mixture was centrifuged at 500 × g for 5 min to remove the unbound EV proteins. The beads were washed with PBS 3 times briefly. The precipitated proteins were subjected to 10% sodium dodecyl sulfate (SDS)-PAGE, and the gel was stained using a silver stain kit (Beyotime, China) according to the manufacturer’s instructions. The gel bands of interest were excised and the proteomic profiles were analyzed and provided by Novogene Corporation Inc.Novogene Co., Ltd. (Beijing, China).

For the immunoprecipitation of fibronectin on EVs, 50 μg GST-N_EV3_ protein and 200 μg EVs derived from differentiated C2C12 cells were incubated in 200 μl PBS at 4°C for 2 h. Then, 0.1% Triton X-100 was added to lyse EVs for 1 h, followed by the addition of GST-beads and incubation overnight at 4°C. Next day, the beads were washed 3 times with PBS and the precipitated proteins were subjected to 8% SDS-PAGE and transferred onto a PVDF membrane. The membrane was probed with anti-fibronectin antibody (1:1000, ImmunoWay, China) to detect the presence of fibronectin in the complex. Anti-Alix (1:200) and anti-CD81 (1:200; Santa Cruz Biotechnology) were used to verify the positive control EVs derived from differentiated C2C12 cells.

### Tissue distribution

To examine the biodistribution of N_EV3_ bound to endogenous circulating EVs, 300 μg mCherry-N_EV3_ or mCherry-N_GFP_ was intravenously injected into *C57BL/6* mice as a single injection. Perfusion was performed 2 h after injection with 50 ml cold PBS to wash out protein in circulation and body-wide tissues were collected. For the N_GFP_-N_EV3_ biodistribution *in vivo*, 100 μg N_GFP_-N_EV3_ was incubated with 100 μg EVs derived from 293T cells at 4°C for 2 h. GFP (200 μg) was administered to *C57BL/6* mice via tail-vein injection, followed by intravenous injection of the N_GFP_-N_EV3_-EVs at 10 min after GFP injection. Tissues were collected 1 h after last administration with perfusion conducted as described above. To examine the tissue distribution of siRNA-DRBD-N_EV3_ loaded onto endogenous circulating EVs, Rhodamine-labeled siRNA (100 μg) was incubated with DRBD-N_EV3_ (1 mg) on ice for 30 min with protection from light. Subsequently, Rhodamine-labeled siRNA or the siRNA-DRBD-N_EV3_ complex was intravenously injected into *C57BL/6* mice. Tissues were collected 30 min after administration with perfusion. To examine the tissue biodistribution of the anti-transferrin receptor (anti-TfR)-protein A-N_EV3_ complex in wild-type *C57BL/6* mice, 500 μg anti-TfR and 100 μg protein A-N_EV3_ were mixed and incubated at 4°C for 2 h. DiR-labeled mouse plasma-derived EVs (500 μg) were intravenously injected to *C57BL/6* mice, immediately followed by intravenous injection of the resulting complex containing anti-TfR and protein A-N_EV3_, separately. Tissues were collected 2 h after administration with perfusion and imaged using the IVIS Spectrum (PE, Waltham, MA, US). For mCherry-N_EV3_, liver tissues were snap-frozen in isopentane precooled with liquid nitrogen for cryosections.

### Protein extraction and Western blot

EV or cell pellets and muscle tissues were lysed in lysis buffer and subjected to 10% SDS-polyacrylamide gel electrophoresis and transferred to PVDF membrane. Membranes were blocked with 5% skimmed milk and incubated with primary antibodies including polyclonal rabbit anti-His (1:2,000), anti-fibronectin (1:1,000; ImmunoWay, China) and anti-α-actinin (1:3000; Sigma, US); and mouse monoclonal antibodies including anti-Alix (1:200), anti-CD63 (1:200), anti-CD81 (1:200; Santa Cruz Biotechnology, US), anti-GAPDH (1:5,000), anti-β-tubulin (1:5,000; Elabscience, China), anti-α-sarcoglycan (1:200), and anti-β-dystroglycan (1:200; Novocastra, UK). Bound primary antibodies were detected by horseradish peroxidase-conjugated goat anti-mouse or goat anti-rabbit IgG (1:5,000; Sigma, US) and the ECL western blot analysis system (Millipore, Billerica, MA, US).

### Immunohistochemistry and histology

Sections of 8 µm were cut from quadriceps and liver and were then examined for mCherry or DiR signals with a confocal fluorescence microscope (Olympus FV1000, Olympus, Japan). The serial muscle sections were stained with a panel of polyclonal and monoclonal antibodies for the detection of DAPC components. Mouse monoclonal antibodies against β-dystroglycan and α-sarcoglycan (1:100, Novocastra, UK), and a rabbit polyclonal antibody against laminin (1:400; Sigma, US) were used according to the manufacturer’s instructions. The M.O.M. blocking kit (Vector Laboratories, Burlingame, CA) was applied for the immunostaining of the DAPC. Mouse and rabbit primary antibodies were detected using Alexa Fluor 594-conjugated goat anti-mouse and goat anti-rabbit IgG secondary antibodies (1:200; Invitrogen, US), respectively. IgG deposition in muscle sections was detected using Alexa Fluor 594-conjugated goat anti-mouse IgG (1:200; Invitrogen, US). Muscle fibrosis was assessed using a Masson’s trichrome staining kit (Solarbio, China) and collagen staining according to the manufacturer’s instructions. A rabbit monoclonal antibody against collagen 1A1 (1:500; ABclonal, China) was used, followed by color detection using a commercial chromogenic detection kit (SZSGB, China). All sections were examined using a Leica upright microscope (Leica Microsystems, Germany). Liver and kidney tissues were fixed overnight in 4% PFA, embedded in paraffin and sectioned at 4 μm. Routine H&E (hematoxylin and eosin) staining was used to examine the overall liver and kidney morphology and assess the level of infiltrating mononuclear cells.

### Clinical biochemistry

Mouse blood was taken from the retro-orbital sinus of mice and allowed to clot at RT for 1 h. Serum was separated by centrifugation at 1,000 × g for 20 min at 4°C and stored at -80°C. Analysis of levels of serum creatine kinase (CK) was performed by the clinical laboratory (Division of Medical Technology, Tianjin Medical University, Tianjin, China). For urinary biochemical analysis, *MRL/lpr* mice were housed individually in metabolic cages for 16 h at the experimental endpoint, with free access to food and water. Urine samples were centrifuged at 3,000 × g for 10 min at 4°C to remove debris, and the resulting supernatants were stored at -80°C until further analysis. Urinary albumin concentrations were measured using a Mouse Albumin ELISA Kit (ICL, US), and urinary creatinine concentrations were determined using a creatinine oxidase assay kit (Nanjing Jiancheng Bioengineering Institute, China) according to the manufacturers’ instructions. The urinary albumin-to-creatinine ratio was calculated by dividing the urinary albumin concentration by the corresponding urinary creatinine concentration and was expressed as milligrams of albumin per gram of creatinine.

### Functional grip strength

Treated and control mice were tested using a commercial grip strength monitor (Chatillon, West Sussex, UK). Briefly, each mouse was held 2 cm from the base of the tail, allowed to grip a protruding metal triangle bar attached to the apparatus with their forepaws and pulled gently until it released its grip. The force exerted was recorded and five sequential tests were carried out for each mouse, averaged at 30 s intervals and averaged. Subsequently, the force readings for force recovery were normalized to body weight.

### Biolayer interferometry

The interaction between N_EV3_ and EVs was tested using an Octet RH16 instrument (Sartorius/ForteBio, Fremont, CA, US) according to the manufacturer’s instructions. The anti-GST biosensors were pre-wetted in PBST buffer containing 0.02% Tween 20 for 10 min. Subsequently, 2 μg GST-N_EV3_ protein was dissolved in 200 μl PBST buffer containing 0.02% Tween 20 for 180 s during the loading step. EVs from 293T cells or from *fibronectin*-knockout 293T cells (2 × 10^9^ particles) were added to 200 μl PBST buffer for 180 s during the association step. Subsequently, the data were analyzed using the Octet Analysis Studio 13.0.

### Molecular docking prediction

The structural complex of human fibronectin (UniProt ID: P02751) and N_EV3_ was predicted using the AlphaFold-Multimer mode on the AlphaFold2 online server (https://alphafold.ebi.ac.uk/). All structural representations and interaction interface illustrations of the predicted models were generated using PyMOL (version 2.5, Schrödinger, LLC).

### Statistical analysis

All data are reported as mean values ±SD. Statistical differences between treatment and control groups were evaluated by Sigma Stat (Systat Software, London, UK). Both parametric and non-parametric analyses were applied as specified in the corresponding figure legends.

## ACKNOWLEDGMENTS

This work was supported by National Natural Science Foundation of China (No. 82320108013, 82571599 and 82630061), National Key R&D Program of China (No. 2024YFA 1802700), Science and Technology Planning Projects of Tianjin Municipality (No. 25ZXZSSS00030), and Tianjin Municipal 15th five-year plan (Tianjin Medical University Talent Project). We thank the Core facility of Research Center of Basic Medical Sciences (Tianjin Medical University) for technical support, specifically the animal imaging and the flow cytometry core facilities.

## AUTHOR CONTRIBUTIONS

H.Y. and G. H. designed the project. G.H., Z.L., Y.H., R.H., Y.L., B.W., B.Wu., T.Z., X.C., J. W. and Y. Z. carried out the experiments. G.H. Z.L. and H.Y. analyzed the data. H.Y., G.H., Y.S. and T.Y. wrote and revised the paper with the input from all authors.

## DECLARATION OF INTERESTS

H.Y., G.H. and L.Z have been granted a patent (ZL202610695954.1). The authors declare no competing interests.

---

**Supplementary Figure 1.**
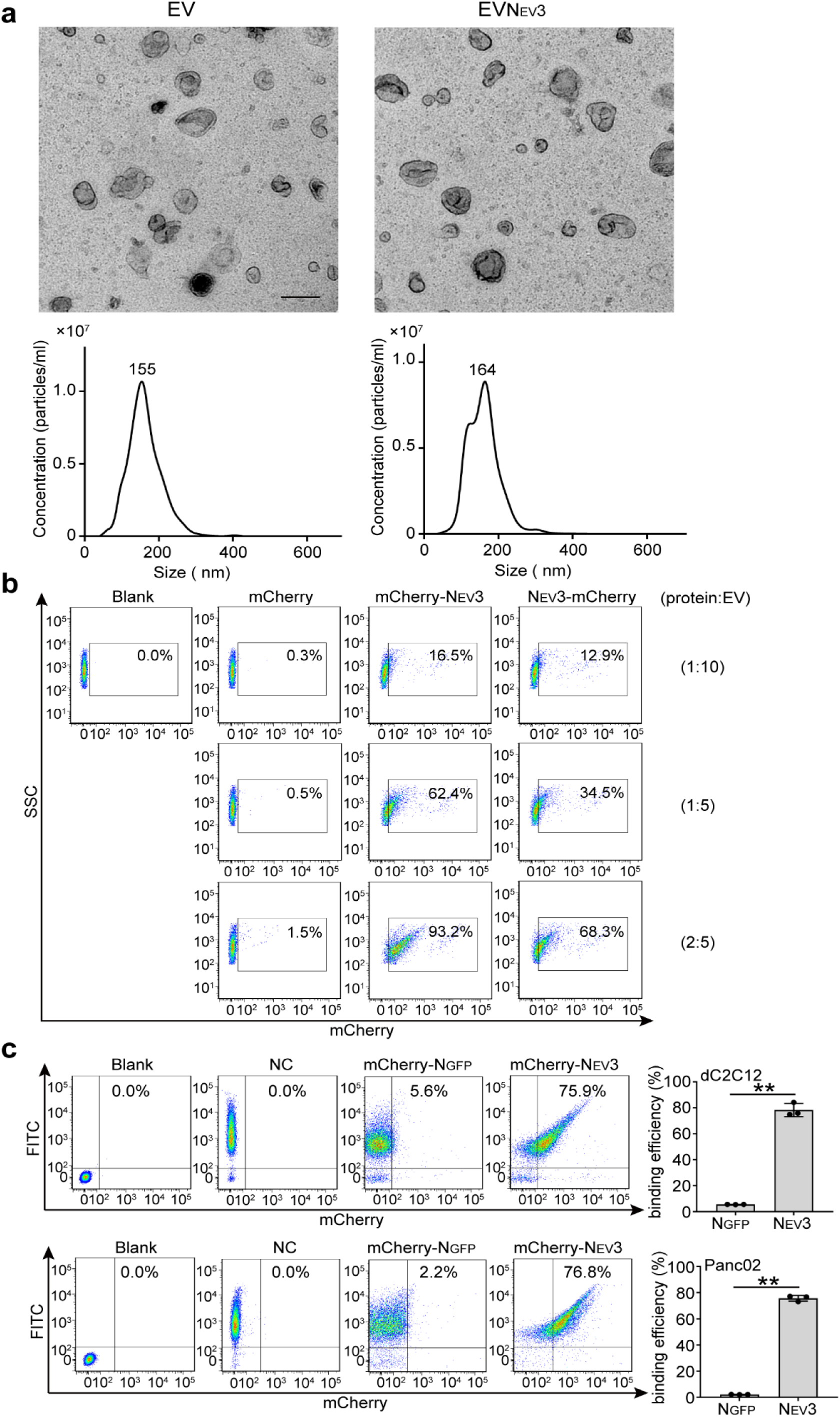
Characterization of differentiated mouse myotube-derived EVs and EVs loaded with N_EV3_, and *in vitro* binding assay. (**a**) Representative transmission electron microscopic (TEM) images and nanoparticle tracking analysis (NTA) of EVs derived from murine myotubes (scale bar=100 nm). (**b**) Representative flow cytometry analysis of the binding of mCherry, mCherry-N_EV3_ or N_EV3_-mCherry to EVs derived from 293T cells at different ratios. Blank refers to EVs alone. (**c**) Representative flow cytometry plots and quantification of the binding of mCherry-N_EV3_ or mCherry-N_GFP_ to EVs derived from mouse myotubes (dC2C12) or Panc02 cells (Panc02) (n=3; **p< 0.01; two-sided unpaired Welch’s t-test). EVs were labeled with WGA-AF488. NC refers to WGA-AF488-labeled EVs. Blank represents unlabeled EVs.

**Supplementary Figure 2.**
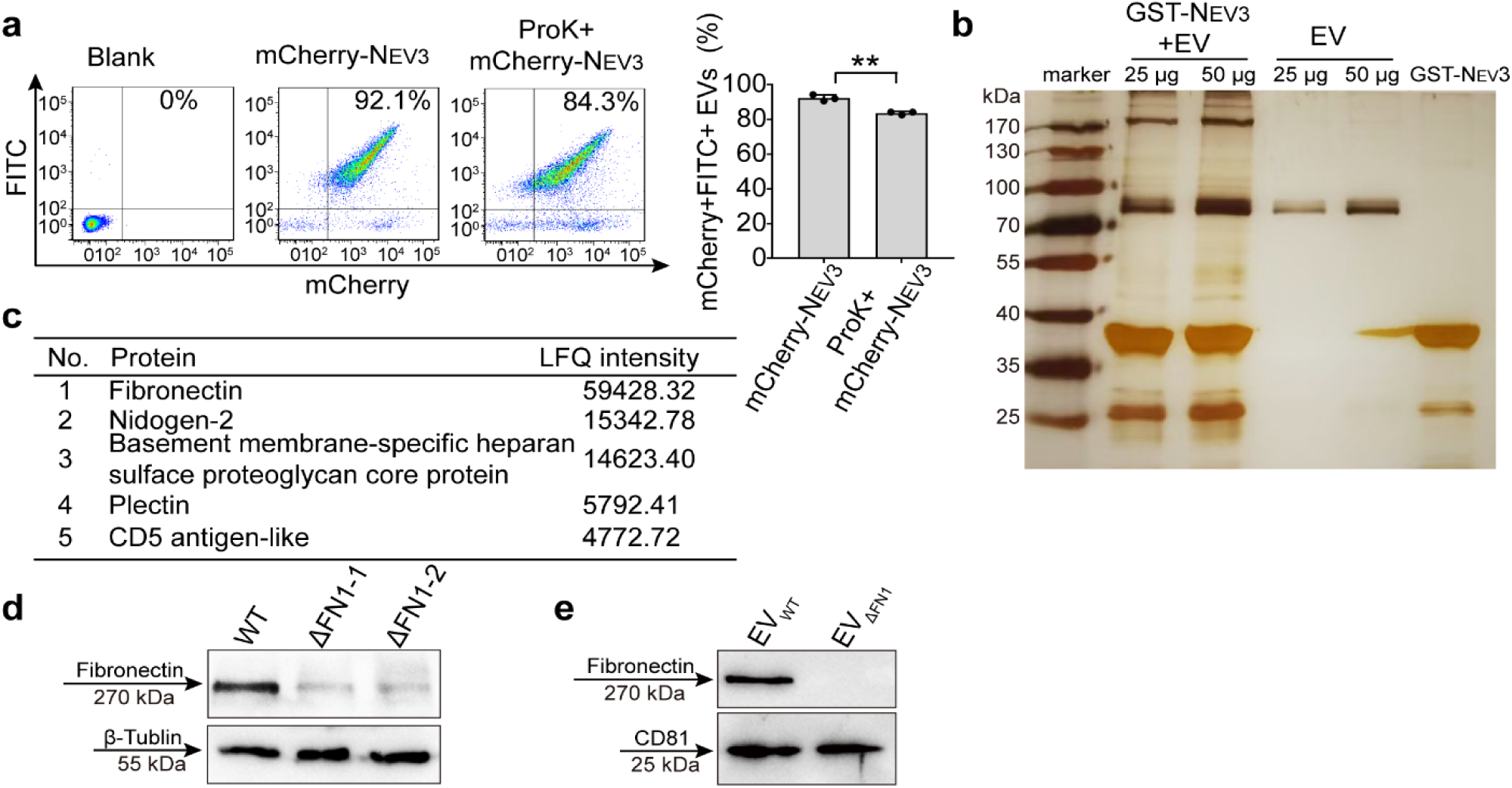
Investigation of N_EV3_ binding partners on EVs. (**a**) Representative flow cytometry plots and quantification of mCherry-positive EVs (n=3; **p < 0.01; two-sided unpaired Welch’s t-test). EVs derived from differentiated mouse myotubes were digested with proteinase K (ProK), followed by incubation with mCherry-N_EV3_. EVs were labeled with WGA-AF488. Blank refers to unlabeled EVs. (**b**) Silver staining of SDS-PAGE gel. Proteins were recovered with GST-N_EV3_-coated beads from EV lysates. EVs were derived from differentiated mouse myotubes. EVs alone and GST-N_EV3_ alone were used as negative controls. (**c**) Table of top candidate proteins identified by mass spectrometry analysis based on label-free quantification (LFQ) intensity. (**d**) Western blot analysis of fibronectin in wild-type 293T cells and two fibronectin-depleted clones (ΔFN1-1 and ΔFN1-2). β tubulin was used as the loading control and a total of 50 μg protein was loaded per lane. (**e**) Western blot analysis of fibronectin in EVs derived from wild-type 293T cells or the fibronectin-depleted clone (ΔFN1-1). CD81 was used as the EV control and a total of 50 μg protein was loaded per lane.

**Supplementary Figure 3.**
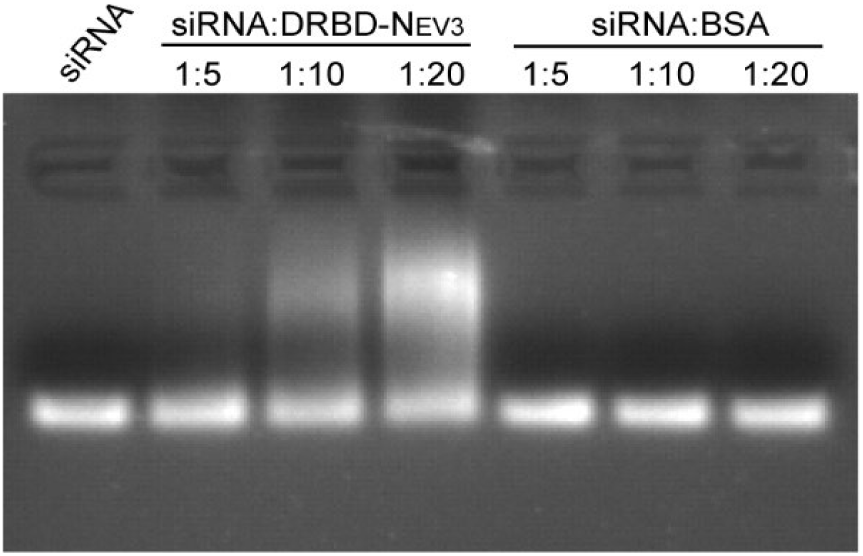
*In vitro* gel migration assay for measuring the binding of DRBD-N_EV3_ to siRNA. Agarose-gel electrophoresis of siRNA after incubation with DRBD-N_EV3_ or bovine serum albumin (BSA) at the indicated siRNA-to-protein mass ratios. Untreated siRNA and BSA were used as negative controls.

**Supplementary Figure 4.**
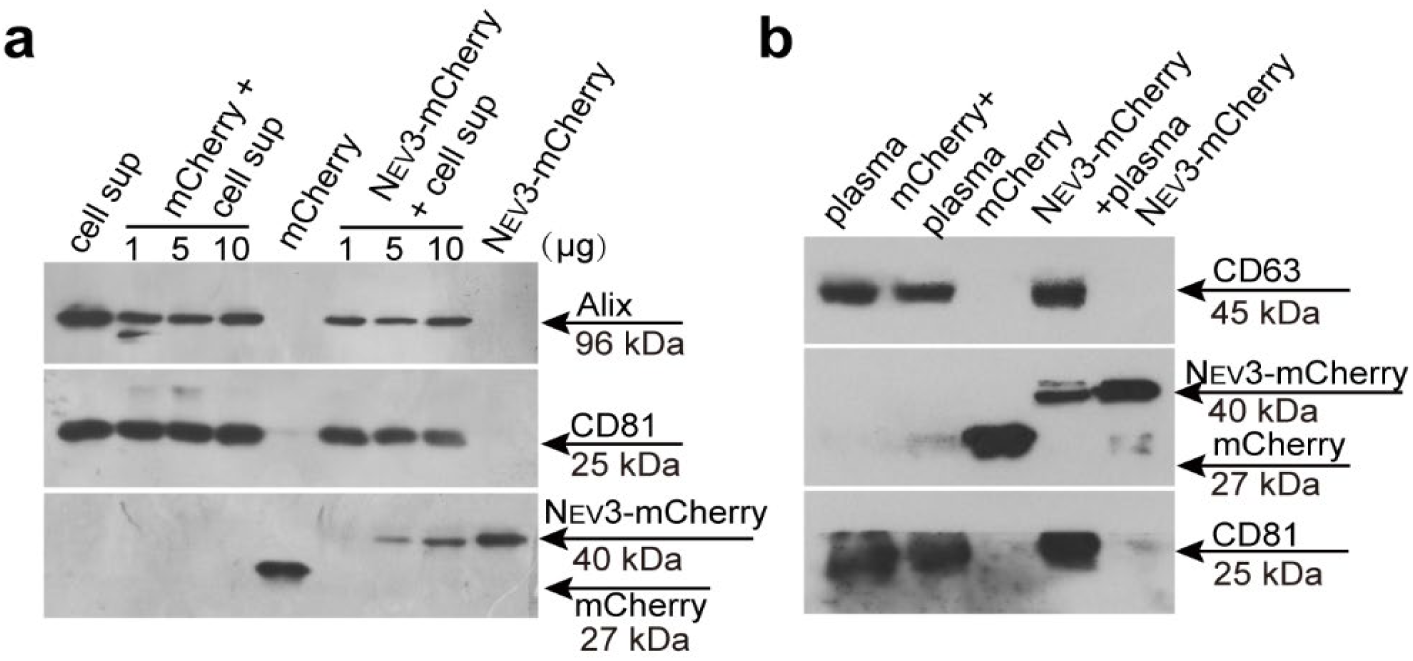
Evaluation of N_EV3_-mediated capture of circulating EVs and protein loading onto EVs. Western blot analysis of mCherry on EVs from cell supernatants (**a**) or from mouse plasma (**b**). Ten μg N_EV3_-mCherry was incubated with cell supernatants from 293T cells or mouse plasma, followed by isolation of EVs by ultracentrifugation. Purified EVs and the proteins alone were used as controls. A total of 50 μg protein was loaded per lane.

**Supplementary Figure 5.**
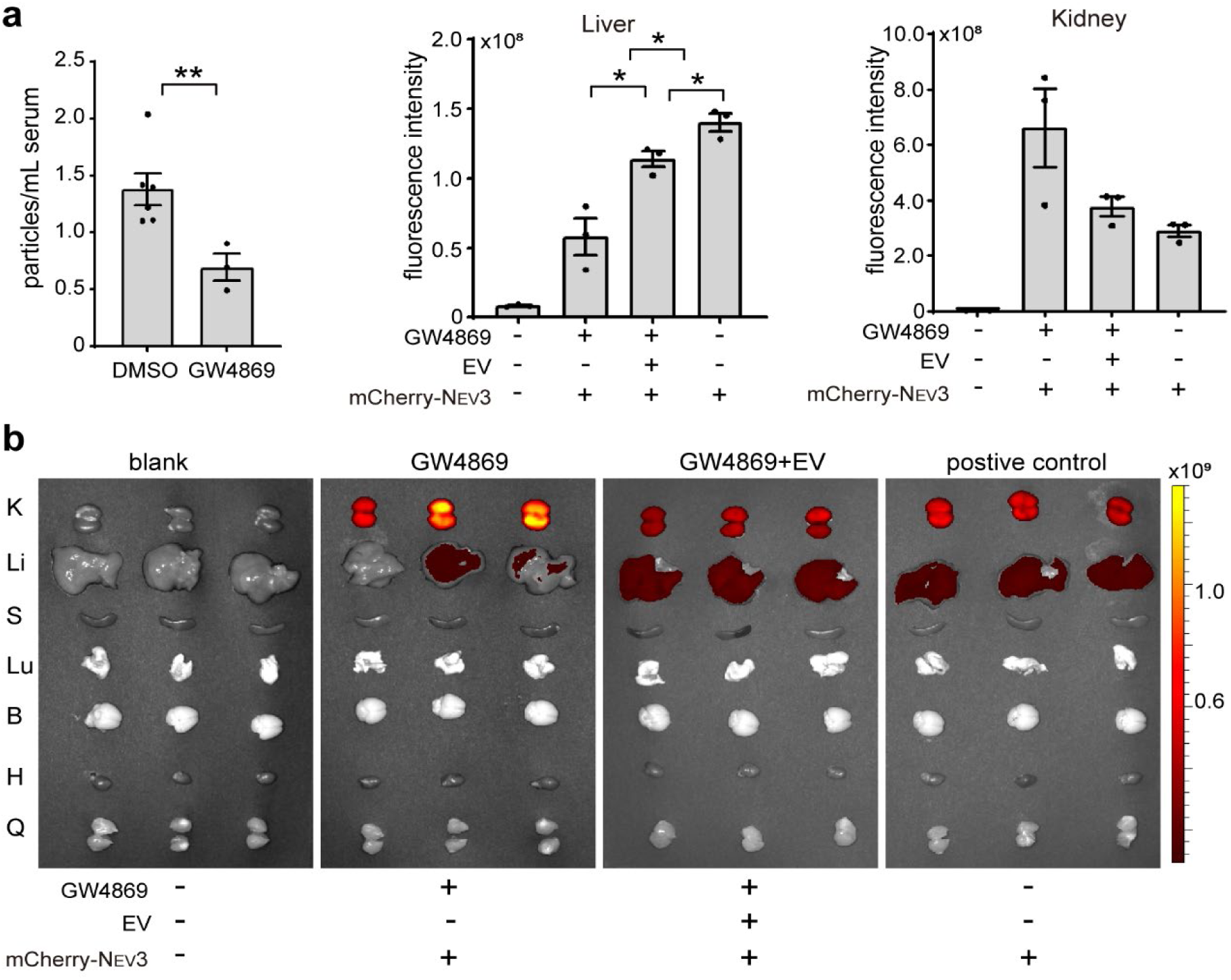
Evaluation of the effect of circulating EVs on N_EV3_’s tissue distribution *in vivo*. (**a**) Nanoparticle tracking analysis of EV concentrations in serum from *C57BL/6* mice treated with DMSO (n=6) or GW4869 (n=3). (**b**) Representative *ex vivo* fluorescence images and quantification of mCherry signals in body-wide tissues from *C57BL/6* mice treated with GW4869 with or without external supplementation of EVs (n=3). mCherry-N_EV3_ was used as a positive control. K, kidney; Li, liver; S, spleen; Lu, lung; B, brain; H, heart; Q, quadriceps. Statistical significance was assessed using Welch’s one-way ANOVA followed by Holm-adjusted pairwise Welch’s t-tests; *p < 0.05; **p< 0.01.

**Supplementary Figure 6.**
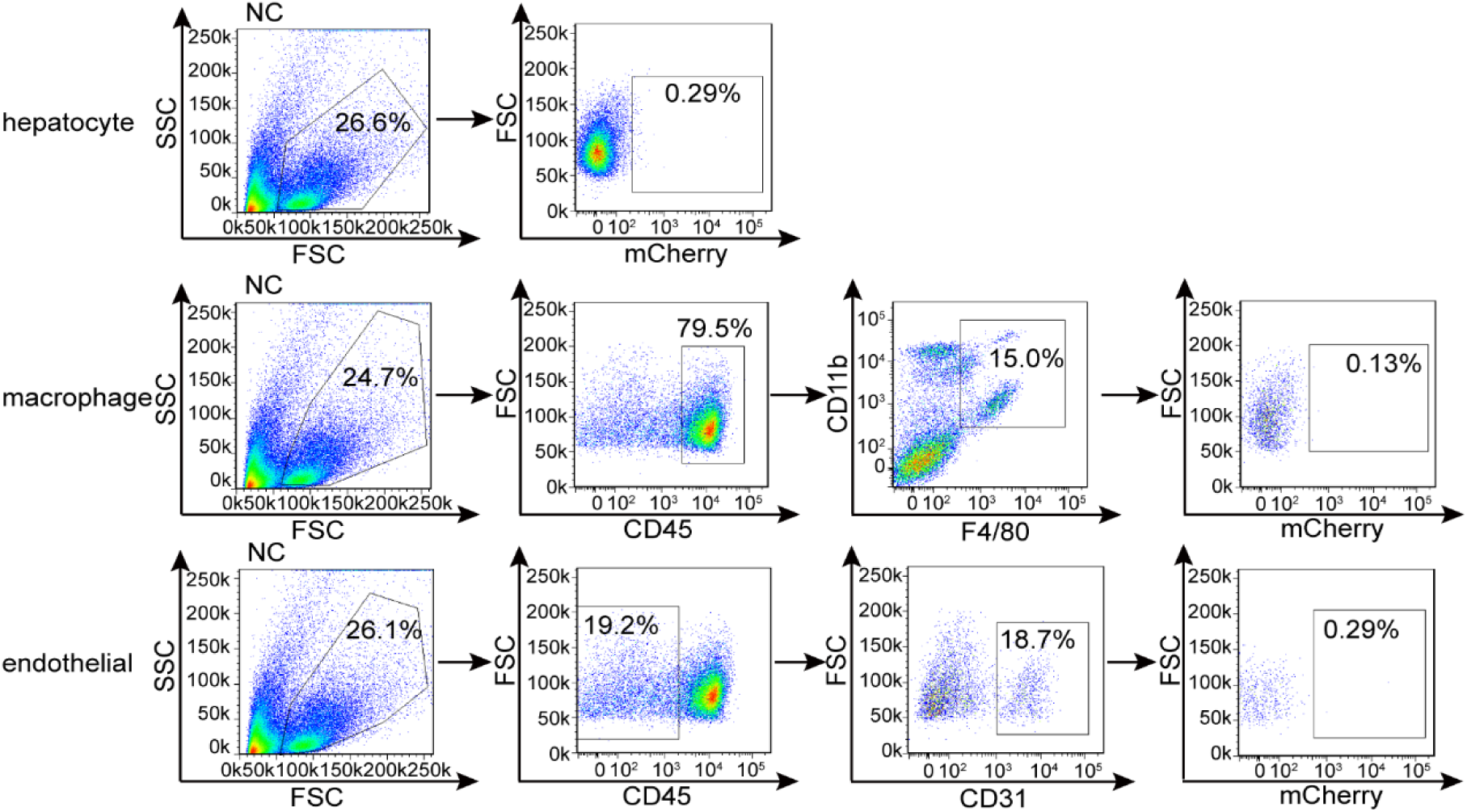
Gating strategy used for flow cytometric analysis of mCherry signals in hepatocytes, macrophages and endothelial cells in the livers of *C57BL/6* mice injected with mCherry-N_EV3_ intravenously.

**Supplementary Figure 7.**
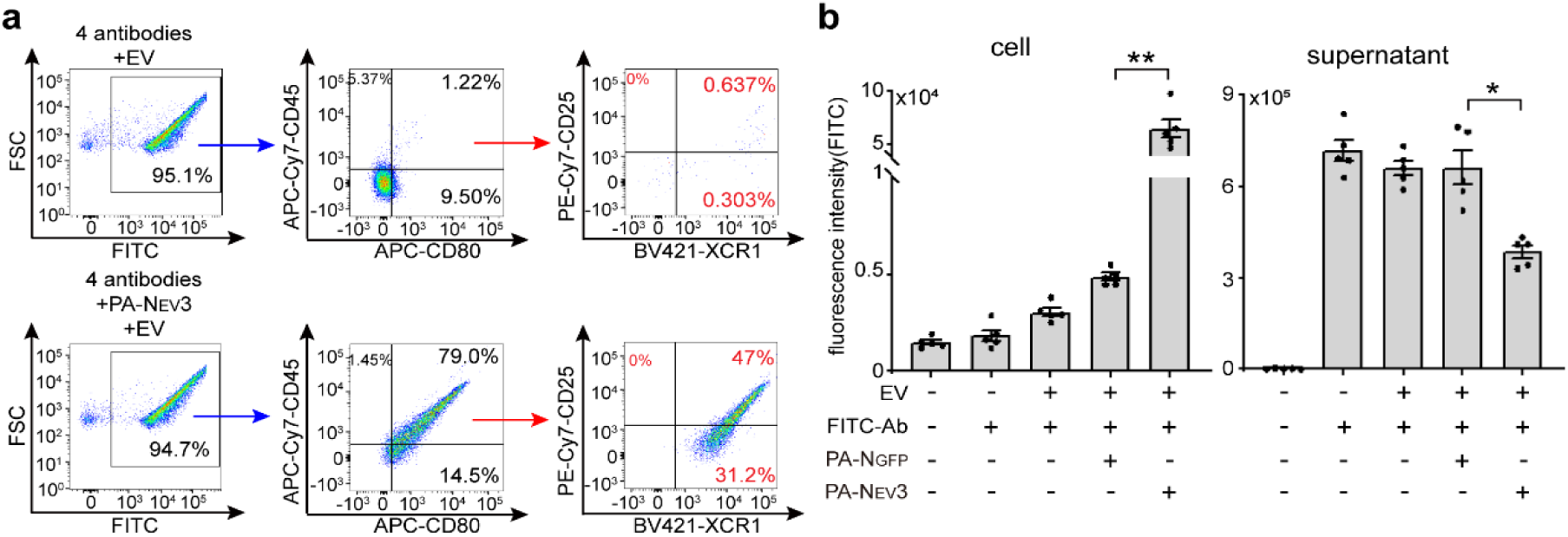
Evaluation of protein A-N_EV3_-mediated antibody loading onto EVs and cellular uptake *in vitro*. (**a**) Gating strategy used to quantify protein A-N_EV3_-mediated simultaneous loading of four fluorescent antibodies, including APC-Cy7-anti-CD45, APC-anti-CD80, BV421-anti-XCR1 and PE-Cy7-anti-CD25, onto EVs derived from 293T cells. (**b**) Quantification of FITC-IgG fluorescence in cells and the corresponding supernatants after separate addition of FITC-IgG, EVs and protein A-N_EV3_ or protein A-N_GFP_ (n = 5; Welch’s one-way ANOVA followed by Holm-adjusted pairwise Welch’s t-tests; *p < 0.05; **p < 0.01). PA refers to protein A.

## Notes

### Competing Interest Statement

The authors have declared no competing interest.

